# Genome-scale community modelling reveals conserved metabolic cross-feedings in epipelagic bacterioplankton communities

**DOI:** 10.1101/2023.06.21.545869

**Authors:** Nils Giordano, Marinna Gaudin, Camille Trottier, Erwan Delage, Charlotte Nef, Chris Bowler, Samuel Chaffron

## Abstract

Marine microorganisms form complex communities of interacting organisms that influence central ecosystem functions in the ocean such as primary production and nutrient cycling. Identifying the mechanisms controlling their assembly and activities is a major challenge in microbial ecology. Here, we integrated *Tara* Oceans meta-omics data to predict genome-scale community interactions within prokaryotic assemblages in the euphotic ocean. A global genome-resolved co-activity network revealed a significant number of inter-lineage associations across large phylogenetic distances. Identified co-active communities included species displaying smaller genomes but encoding a higher potential for quorum sensing, biofilm formation, and secondary metabolism. Community metabolic modelling revealed a higher potential for interaction within co-active communities and pointed towards conserved metabolic cross-feedings, in particular of specific amino acids and group B vitamins. Our integrated ecological and metabolic modelling approach indicates genome streamlining and metabolic auxotrophies as central joint mechanisms shaping bacterioplankton community assembly in the surface global ocean.

## Main

Marine microbes constantly interact among each other and with their environment, forming complex and dynamic networks. These communities and their interactions play crucial ecological and biogeochemical roles on our planet, forming the basis of the marine food web, sustaining biogeochemical cycles in the ocean, and regulating climate^1^. Complex networks of trophic interactions, mediated through metabolic cross-feeding and ecological successions, can influence the nature of microbial interactions (e.g., mutualism or competition), in space and time, and thus significantly shape microbial community assembly^2^. Expanding our understanding of microbial trophic interactions is fundamental given their capacity to modulate ecological niches^3^, constrain microbial biogeography^4^, drive microbial diversification^5^, and modulate the eco-evolutionary dynamics of microbial communities^6^. Because most microbes are difficult to isolate and cultivate in lab-controlled environments^7^, and given the large diversity of molecules that can be excreted into the environment (e.g., waste metabolites, secondary metabolites, exoenzymes, siderophores), we are just starting to grasp the complexity and diversity of microbial interactions and cross-feeding relationships existing in nature^8^. In particular, we lack a mechanistic understanding of metabolic auxotrophy and its role in constraining marine microbial community composition and assembly^9^.

While species co-occurrence networks are useful tools to model the large-scale structure of microbial communities^10^ and to resolve biome-specific ecological associations^11^, these approaches are inherently limited since correlation metrics do not provide evidence for direct biotic interactions, and do not allow to disentangle true biotic interactions from environmental preferences (niche overlap)^12^. Thus, we still lack a comprehensive and mechanistic understanding of biotic and abiotic interactions shaping community assembly of microbial communities. Ecosystem modelling approaches are therefore needed to capture and predict emergent properties resulting from complex interactions within microbial communities, such as resilience, niche space, and biogeography, that shape microbial communities and ecosystems^13^. Recent experimental work has demonstrated the significant impact of underlying cross-feeding metabolic networks in shaping community assembly^14^ and ecological successions^15^ in synthetic microbial communities. Using microbial community assembly experiments in soil, coupled with a simple resource-partitioning model, functional convergence was shown to be mainly driven by emergent metabolic self-organization, while taxonomic divergence seemed to arise from multi-stability in population dynamics^14^. In another system, coculture experiments of a marine microbial community able to degrade chitin demonstrated the hierarchical preferences for specific substrates, underlining the sequential colonization of metabolically distinct groups, and identifying hierarchical cross-feedings shaping the dynamics of community assembly^15^.

Recent large-scale environmental surveys of marine microbial ecosystems (e.g., *Tara* Oceans^16^, Malaspina^17^, Bio-GO-SHIP^18^, BioGEOTRACES^19^) have generated large volumes of metagenomics data that enable the reconstruction of genomes from uncultivated species referred to as Metagenome-Assembled Genomes (MAGs)^20,21^. Together with whole genome sequences (WGS) from cultured organisms and single amplified genomes (SAGs) from single cell isolates, these resources have been used to expand our knowledge of microbial diversity in the ocean, but have also demonstrated that a large fraction of the diversity remains to be explored^22,23^. In this context, genome-resolved metagenomics provides an opportunity to enrich co-occurrence signals with genetic information from genomes and functional information from genome-scale metabolic models. Integrating this knowledge into association networks can inform us about the functional self-organisation of microbial communities^24^, contribute to our understanding of species interactions mechanics, and identify general ecological laws that structure microbial communities. While community metabolic modelling approaches have recently been applied to study the self-organisation of microbial ecosystems^25^ and to gain insights into molecular mechanisms of interactions in soil^26^, wastewater^27^, and gut microbiome communities^28^, few studies so far have focused on the modelling of marine plankton ecosystems^15,29^, and were limited to specific single communities.

Here, we describe an integrated ecological and metabolic modelling approach with the goal to delineate metabolically cohesive consortia underlying genes-to-community assembly and ecosystem functioning at global scale^30^. We combined co-activity ecological information inferred from meta-omics with community metabolic simulations using genome-scale metabolic models to uncover putative biotic interactions mediated by metabolic cross-feedings among marine prokaryotic genomes. Through a multi-omic approach integrating *Tara* Oceans metagenomic and metatranscriptomic datasets, we inferred a global ocean genome-resolved ecological network from whole-genome transcriptomic activities. We used general genomic scaling laws^31^ as a framework to characterise the functional content of co-active environmental genomes, and identified functional gene categories likely driving metabolic dependencies. We then reconstructed genome-scale metabolic models and uncovered putative cross-feeding interactions within co-active consortia through the use of community-level metabolic modelling.

## Results and discussion

### Genomic scaling laws reveal features of uncultivated marine prokaryotic genomes

To build a comprehensive catalogue of marine prokaryotic genomes, we collected and assembled public whole-genome sequences (WGS) from marine prokaryote isolates^32^, single-amplified genomes^22^ (SAGs), as well as previously reconstructed MAGs^21^. This novel integrated marine prokaryotic genome database counted 7,658 non-redundant species-level representative genomes (delineated by a 95% ANI threshold over 60% of genome length, see methods). Herein, we only considered genomes meeting sufficient quality standards (n=5,678) as defined by the Genomic Standards Consortium^33^, that is High-Quality (HQ) MAGs (>90% complete with less than 5% contamination) as well as Medium-High-Quality (MHQ) MAGs (>75% complete with less than 10% contamination). HQ and MHQ MAGs were not significantly different from WGS genomes in terms of gene density (**Supplementary Table 1**). A phylogeny of these genomes was established using domain-specific marker genes of the Genome Taxonomy Database (GTDB)^34^, highlighting a total of 107 phyla (with unclassified) including highly represented phyla in marine environments, such as Proteobacteria, Bacteroidetes, Actinobacteria, and Cyanobacteria^35^ (**Fig. 1a**).

**Figure 1:**
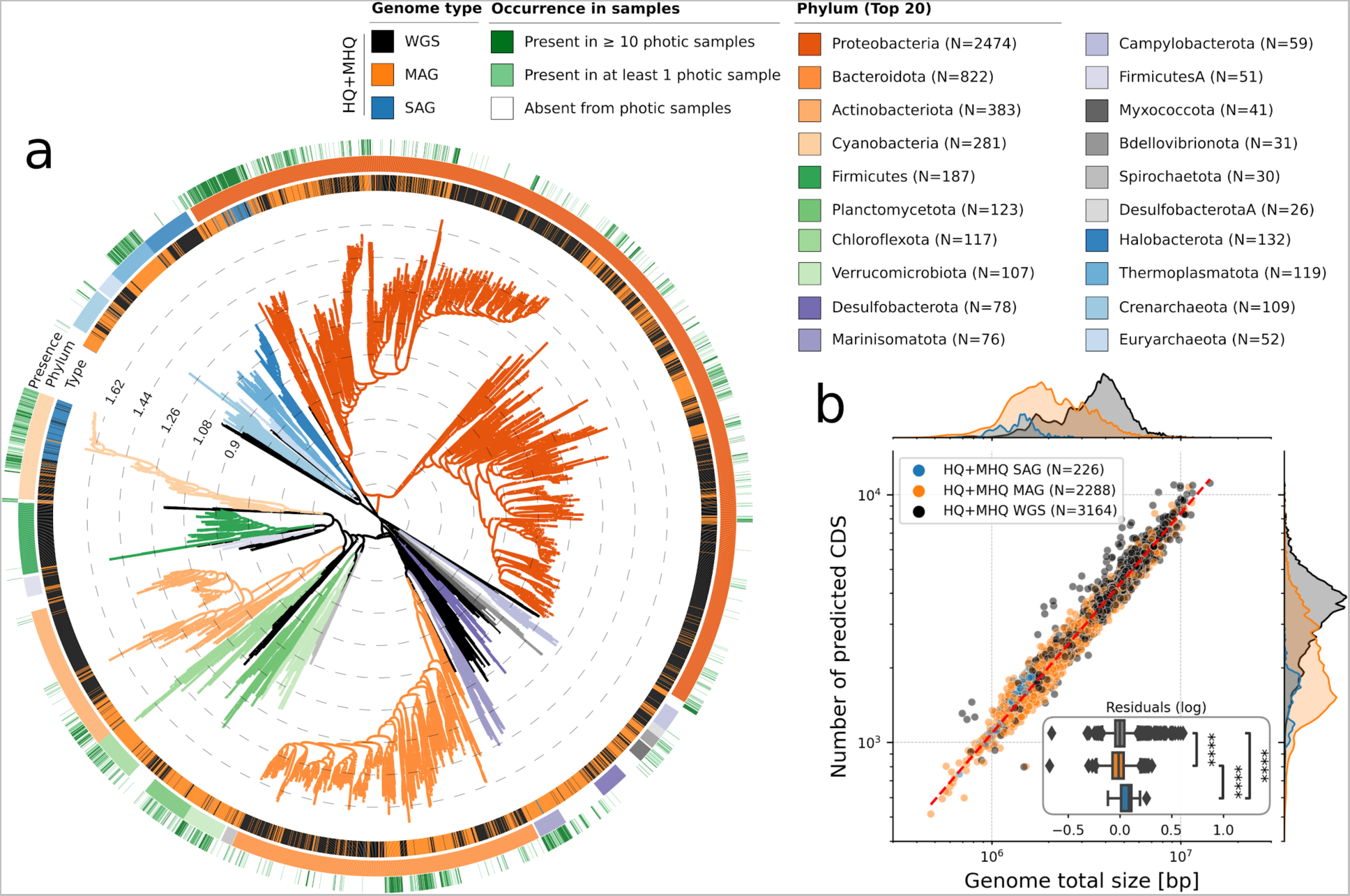
A database of marine bacterial and archaeal genomes from isolates and uncultivated genomes reconstructed from marine metagenomes. **a,** Phylogenetic tree of the database of marine genomes (N=7,658) dereplicated at species level (95% Average Nucleotide Identity or ANI). Reference genomes (WGS) were obtained from MarRef, MarDB, and aquatic progenomes, while Metagenome-Assembled Genomes (MAGs) and Single-Amplified Genomes (SAGs) were also obtained from different studies (see methods). A total of 107 phyla (including unclassified) were detected (the top 20 most represented phyla are highlighted). **b,** A comparison of genome size and number of predicted CDS revealed that a genome scaling law is conserved for High and Medium-High Quality (HQ and MHQ) genomes (completeness ≥ 75% and contamination ≤ 5%), and that MAGs overall displayed significantly smaller genomes (*P*=8.14 × 10^−289^, Mann Whitney U test on log-transformed distributions).

Within prokaryotic genomes, the number of genes in most high-level functional categories has been shown to scale as a power-law to the total number of genes in a genome^36^. A potential explanation for these observed scaling laws among microbial genomes is a conserved average duplication rates for the evolutionary process within each functional category. In addition, these genomic scaling laws have been shown to be conserved across microbial clades and lifestyles, supporting the observation that they are universally shared by all prokaryotes^31^. However, these genomic scaling laws have never been investigated within uncultured genomes so far. Here, we thus revisited this universal law for environmental marine genomes (MAGs and SAGs). To ensure a sound and fair comparison between WGS and environmental genomes, we limited our analysis to HQ and MHQ genomes, which were of equivalently high-quality and also having a similar gene density as compared to WGS (**Extended Data Fig. 1b**). We showed that HQ and MHQ genomes did actually fit the same law as WGS genomes (**Fig. 1b**). This analysis also revealed that HQ/MHQ MAGs and were systematically smaller in genome size and number of predicted CDS as compared with WGS genomes. This observation is coherent with the assumption that naturally occurring marine genomes have likely adapted to oligotrophic surface ocean specific lifestyles through genome streamlining^37^. Investigating the genomic scaling laws for high-level functional categories (see methods), we showed that this adaptation has differentially impacted specific functions within uncultivated genomes (MAGs and SAGs), with an increase potential for xenobiotic degradation, terpenoid and polyketide metabolism, as well as lipid metabolism, but a decrease potential to synthesize cofactors and vitamins (**Extended Data Fig. 2 and Supplementary Table 2**). This decreased metabolic potential for cofactors and vitamins in environmental genomes likely reflects the importance of syntrophic metabolism, such as metabolism of essential enzyme cofactors^38^, and associated bacterial traits for microbial interactions^39^, to sustain microbial life in the surface ocean that is largely depleted in B vitamins^40^.

### Abiotic factors shaping genome community composition and activity

Next, we mapped *Tara* Oceans metagenomics and metatranscriptomics sequencing reads from surface (SRF) and deep chlorophyl maximum (DCM) samples (N=118) onto our genome collection (see methods) to generate a comprehensive global ocean abundance and expression profiling of microbial communities in relationship with abiotic environmental factors (see **Supplementary Table 3**). Average mapping rates were 16.0% and 12.3% for metagenomes and metatranscriptomes, respectively (**Fig. 2a and Extended Data Fig. 3**). Using the same *Tara* Oceans dataset, gene and transcript abundances have previously been shown to be highly correlated^41^. Here, we observed an overall relatively good concordance between genome-wide abundance and expression (Spearman rho=0.68, P=0)), albeit a number of genomes displayed lower genome-wide expression levels (**Fig. 2b**), highlighting the complementary information brought by genome expression signals computed here. Thus, this observation prompted us to compute genome-wide activities, integrating abundance and expression levels at the genome scale (see methods). Principal Coordinates Analyses (**Fig. 2c**) did not reveal a clear structuration of community genome assemblages and activities by ocean basin, but allowed us to identify abiotic factors driving community composition and activity. Genome community composition was mainly driven by temperature, pH, and Photosynthetically Available Radiation (PAR), while genome community activity was mainly driven by temperature, phosphate (PO_4_) and iron concentrations (see methods and **Supplementary Table 4**).

**Figure 2:**
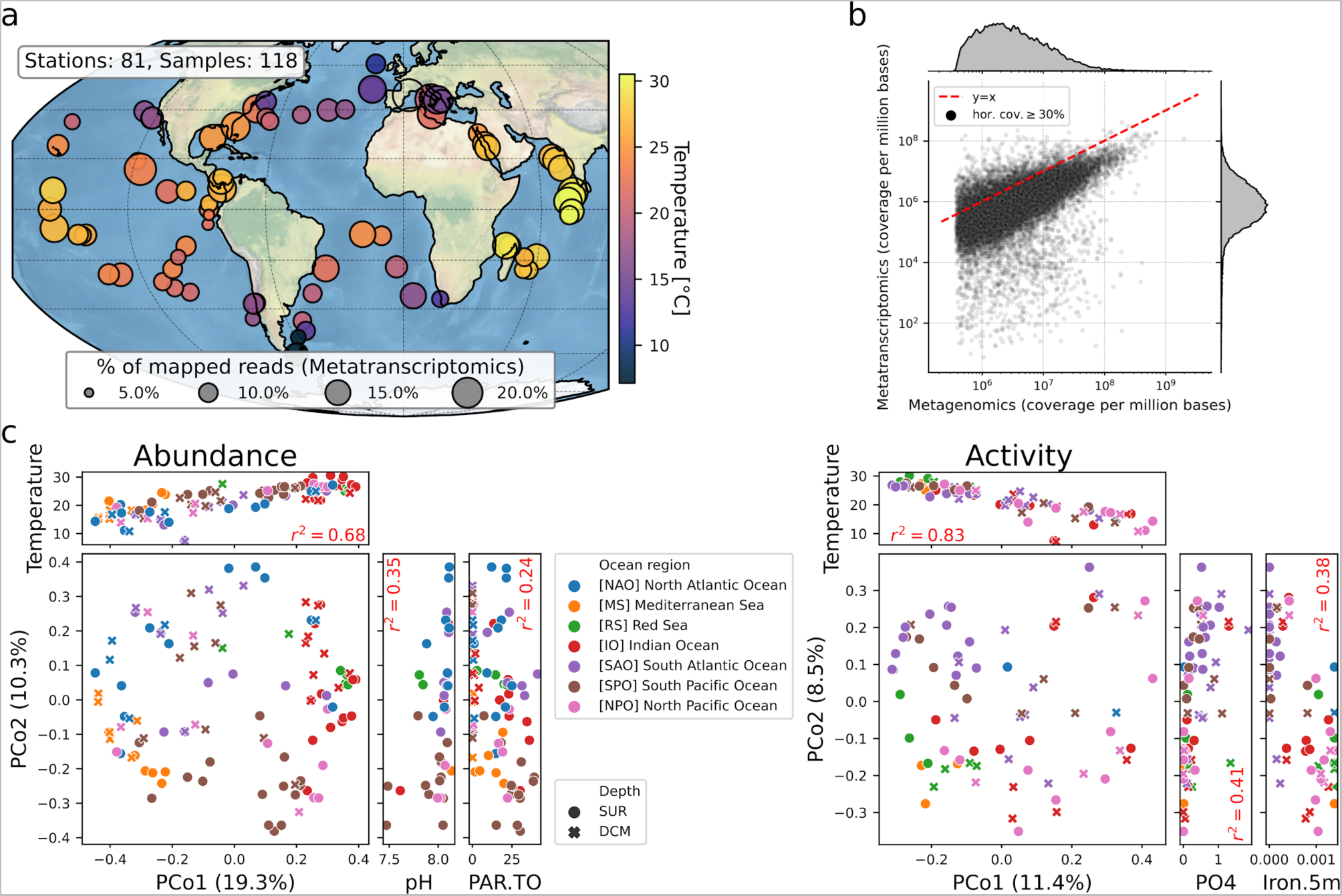
Genome-wide abundance and activity profiling of marine prokaryotic genomes in the global surface ocean. **a,** World map of *Tara* Oceans sampling stations (N=81) for which euphotic (SRF and DCM) metatranscriptomes are available for a prokaryote-enriched size fraction (0.22-3 μm). The percentage of mapped RNA reads are depicted for each euphotic sample (N=118). **b,** Genome-wide abundance and expression were significantly associated (Spearman rho=0.68, P=0), albeit a number of genomes display lower expression levels. **c,** Principal Coordinates Analyses (PCoA) for genome community abundances and activities. Genome community abundance and activity (PCo1) are significantly associated with temperature. Community abundance (PCo2) is also associated with pH and Photosynthetically Available Radiation (PAR), while community activity is associated with PO_4_ and iron concentrations.

Temperature has previously been shown to be one of the main factors constraining epipelagic bacterioplankton community composition^35^, which is confirmed here for both genome-wide community abundance and activity. The effect of (small) pH changes on marine microbial communities has mainly been shown experimentally^42,43^, but often not considering the natural variability of pH in the surface ocean^44^. Other studies have reported minor effects of acidification on the productivity of natural picocyanobacteria assemblages^45^. Here, the observed association between genome community composition and pH could partly be explained by seasonal variability encountered during global sampling. While genome community activity was principally associated to temperature, distinct environmental factors, namely PO_4_ and iron concentrations, were also significantly associated to community activity. This observation emphasises the major role of nutrients and/or cofactors (co-)limitations in structuring global ocean microbial activity^46,47^.

### Biotic drivers of genome activity community structure

While abiotic factors are known to be significant drivers of microbial community structures in the ocean, biotic factors (such as competition, parasitism, or mutualism) are expected to play an equally important role^48^, though the latter are more difficult to study in natural communities. Microbial association networks are useful abstractions that represent potential biotic interactions and capture emergent properties (e.g., connectivity, functional redundancy) that result from these putative interactions^49^. But so far, most studies have been limited to the organismal level by predicting these ecological associations using taxonomic marker genes (e.g., 16S and 18S rRNA genes). Integrating genomic information into association networks can be particularly useful to draw and test hypotheses about the functional self-organisation of microbial communities^24^. Here, we went beyond by inferring a global ocean association network from genome activities that were inferred by integrating genome-wide abundance and transcript levels (here activity refers to a genome-wide ratio between transcript and genomic vertical coverages, see methods for details). We make the general assumption that a co-activity signal is a better proxy to capture biotic interactions as compared to co-abundance, given the latter is an integration of all past metabolic activities that cannot identify microbial cells that were actually transcriptionally active at sampling time. In other words, we expect genome-wide co-activity (integrating abundance and transcript levels) to be more sensitive as it inherently has a better time-resolution when looking for microbial interactions.

We inferred a genome-resolved co-activity network using the dedicated probabilistic learning algorithm FlashWeave (FW, see methods) that can efficiently detect and remove undirect associations among features^50^. This genome-resolved co-activity network was significantly different than the corresponding genome-resolved co-abundance network, with a higher number of edges in co-activity, and only a small fraction of shared edges (3%) (**Extended Data Fig. 4**). This strong difference between both networks can reflect the distinct information carried out by abundance and activity profiles, but can also be partially explained by the heuristics-based inference of direct associations as implemented in FW. The co-activity network revealed a larger number of significant positive associations across large phylogenetic distances (PD), while negative associations were mainly observed between phylogenetically distant genomes (**Fig. 3a**). It also revealed two distinct types of positive associations: relative phylogenetically close associations (0 < PD < 1) that likely reflected niche overlap, and phylogenetically distant associations (PD >= 1) likely reflecting a higher potential for cross-feeding interactions^51^. As previously reported for co-existing genomes across various biomes^24^, co-active genomes tended to be more functionally related than expected at random (Mann-Whitney U-test with Bonferroni correction, P=1.187×10^−25^). This observation may reflect the impact of ecological preferences or niche overlap on evolution, that could be explained by adaptation to a same niche and/or by potential higher rates of horizontal gene transfer (HGT) in specific biomes^52^. Marine co-active genomes also tended to be smaller in size as compared to detected but non-co-active genomes, although displaying similar gene densities as assessed by genomic scaling laws (**Extended Data Fig. 5**), and despite the fact that most abundant and active genomes actually corresponded to MAGs overall smaller in size (**Fig. 1**). In addition, comparative genomics analyses based on scaling laws allowed us to take into account genome size (see methods) and revealed that co-active genomes displayed (in proportion) a higher metabolic potential for lipid, carbohydrate, and amino acid metabolism (**Extended Data Fig. 6 and Supplementary Table 5**), but also for terpenoids and polyketides, quorum-sensing and biofilm formation, as well as for secondary metabolite biosynthesis (**Fig. 3b-d**). Overall, these enriched genomic potentials in co-active genomes point towards key metabolic functions for energy harvest and storage (i.e., lipid, carbohydrate and amino-acids metabolism), likely key in nutrient-limited regions of the global ocean^47^. But they also underline key genomic enriched potential (i.e., antimicrobials and quorum-sensing) of marine genomes likely prone to a wide diversity of biotic interactions^39^.

**Figure 3:**
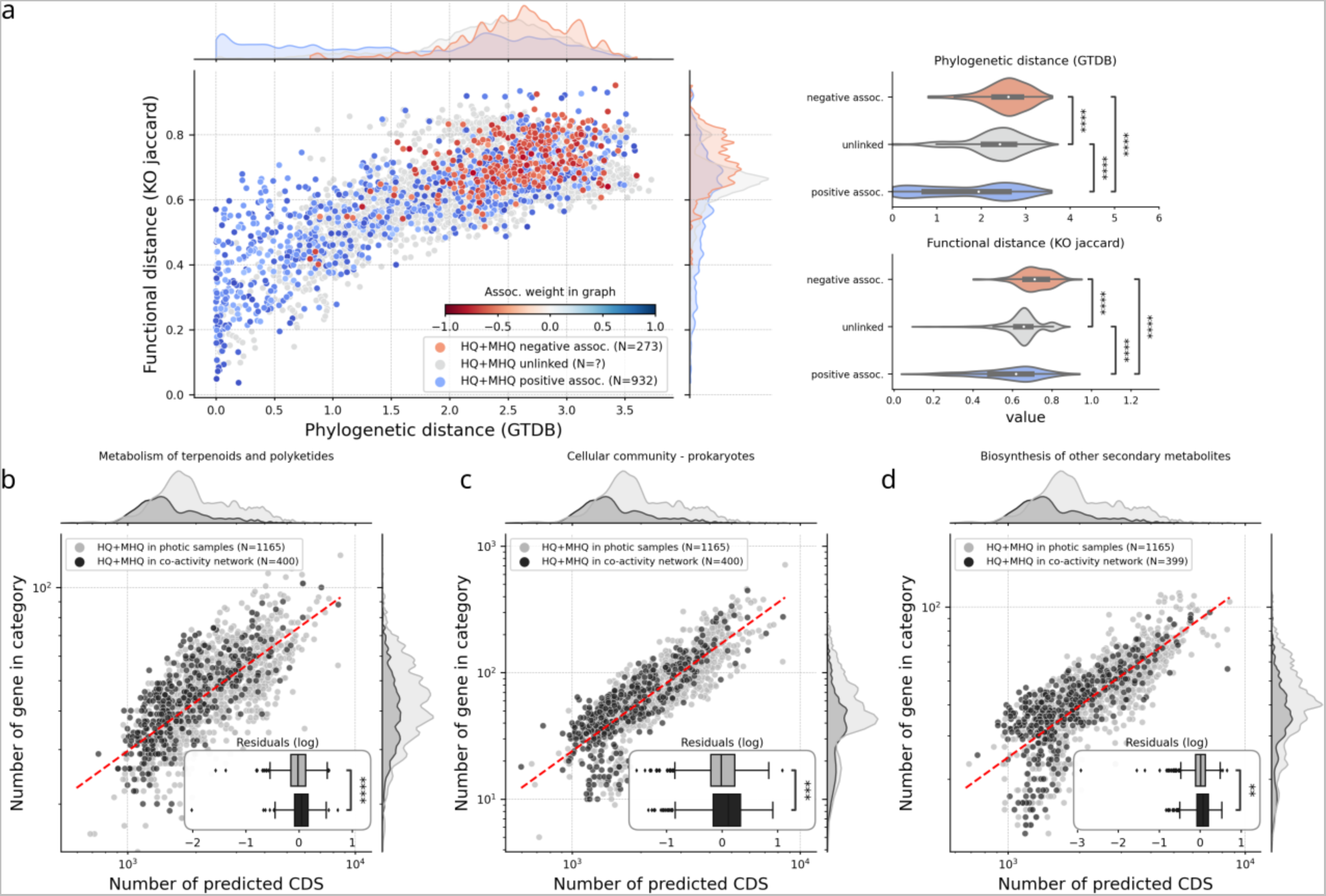
A genome-resolved co-activity network reveals biotic factors shaping marine prokaryotic community structure. **a,** A genome-resolved co-activity network was inferred from genome-wide activities in euphotic samples, and revealed a larger number of significant positive associations between genomes across large phylogenetic distances. Based on gene presence/absence using Jaccard distances between genomes as a proxy for functional distance (using KEGG or eggNOG functional hierarchies), co-active genomes were functionally closer than expected at random (P=1.187×10^−25^). **b,** Scaling laws in the functional content of genomes highlighted large metabolic categories enriched in co-active genomes versus genomes detected as active in samples. Notably, co-active genomes displayed a higher functional potential for terpenoid and polyketide metabolism (P=1.46×10^−7^), for cellular community metabolism (quorum-sensing and biofilm formation, P=4.00×10^−4^), and for the biosynthesis of other secondary metabolites (P=4.73×10^−9^). See Supplementary Table 5 for a complete list of functions enriched in co-active genomes.

### Higher metabolic interaction potential in co-active bacterioplankton communities

To go beyond correlation-based and enrichment analyses and move towards a mechanistic understanding of marine microbial community functioning, we sought to model the community metabolism of co-active marine microbial genomes. To do this, we first reconstructed genome-scale metabolic models for each MHQ and HQ genome (WGS or MAGs) using CarveMe^53^ and quality checked them using MEMOTE^54^ (**Supplementary Materials**). We then used Species Metabolic Coupling Analysis (SMETANA); a constraint-based technique commonly applied for modelling interspecies dependencies in microbial communities^55^. Here, SMETANA was used to compute several interaction scores (global or local) to predict metabolic interaction potential and reveal metabolic exchanges and cross-feedings within delineated communities of co-active genomes. Notably, the Metabolic Resource Overlap (MRO) quantified how much species in a given community compete for the same metabolites, and the Metabolic Interaction Potential (MIP) quantified how many metabolites community species can share to decrease their dependency on external resources. Here, we analysed co-active genome communities identified by clustering the global co-activity network using the Markov clustering algorithm (see methods).

Overall, we observed a negative association between the MRO score and the mean community phylogenetic distance (Pearson R^2^=0.31, P=4.16×10^−8^, **Extended Data Fig. 7**), showing that, as expected, phylogenetically closer co-active genome communities tended to display a higher metabolic resource overlap, and thus a higher potential for competition. Co-active genome communities also displayed an overall lower MIP score as compared with random communities (Mann-Whitney U test, P=1.45×10^−17^, **Extended Data Fig. 8a**). Nevertheless, both global (MIP) and detailed (SMETANA sum) scores of metabolic interactions are significantly driven by the size of communities under consideration (**Extended Data Fig. 8b**), which we thus normalised by community size, as previously done and reported^55^. Following this normalisation and despite overall higher MRO scores and mean community phylogenetic distance, co-active genome communities displayed a higher potential for metabolic interactions as compared with randomly assembled communities (**Fig. 4a**). These results show that metabolic cross-feeding interactions can occur across a large spectrum of phylogenetic and functional distances, suggesting that metabolic dissimilarity is one among other factors determining the establishment of cross-feeding interactions among bacteria^51^.

**Figure 4:**
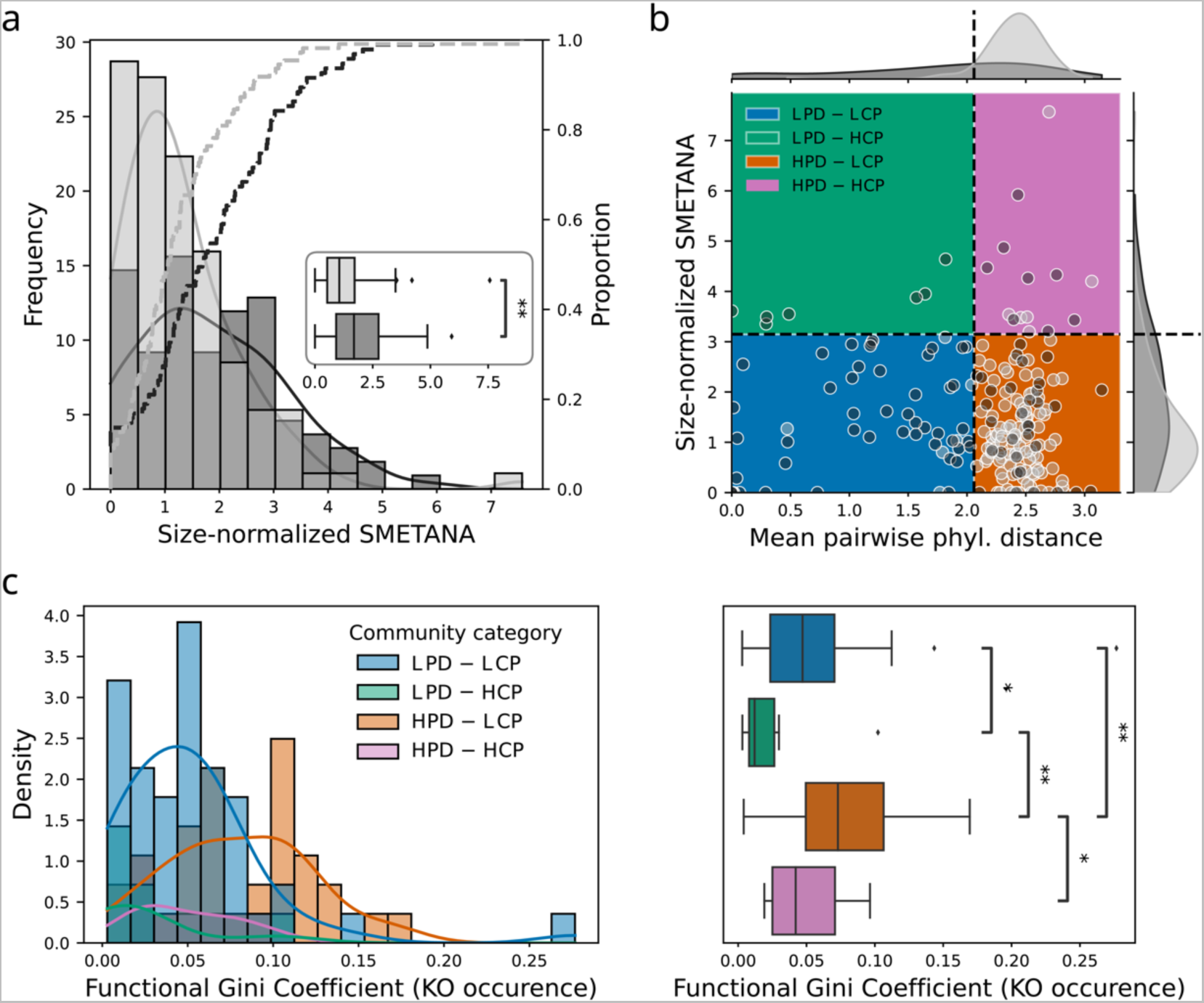
Community-wide metabolic modelling reveals a higher metabolic interaction potential within marine prokaryotic communities. **a,** Microbial communities were delineated on the global co-active genome network using the MCL graph clustering algorithm (see methods). Community metabolic modelling was performed using SMETANA on these communities (dark grey, frequencies as bars and proportions as dashed line) and compared to random communities (light grey). Co-active communities (dark grey) overall displayed a significantly higher metabolic interaction potential (SMETANA) score as compared with random communities (Mann-Whitney U test two-sided, P=1.09×10^−3^). **b,** Distinct metabolic interactions community types were identified within co-active marine prokaryotic communities (black points) and differentiated from random communities (grey points), the latter largely displaying an overall higher mean phylogenetic distance and lower metabolic cross-feeding potential score (HPD-LCP, orange quadrant): i) Communities with overall low mean phylogenetic distance and low metabolic cross-feeding potential score (LPD-LCP, blue quadrant), ii) communities with overall low mean phylogenetic distance and high metabolic cross-feeding potential score (LPD-HCP, green quadrant), and iii) communities with overall high mean phylogenetic distance and high metabolic cross-feeding potential score (HPD-HCP, pink quadrant). **c,** HPD communities (orange and pink) were more dissimilar to respective LPD communities (blue and green) according to their functional Gini coefficient inferred from KEGG metabolism KO genes occurrence profiles (Mann-Whitney U test two-sided with Benjamini-Hochberg correction, LPD-LCP vs. LPD-HCP P=4.88×10^−2^, LPD-HCP vs. HPD-LCP P=2.77×10^−3^, HPD-LCP vs. HPD–HCP P= 4.89×10^−2^, LPD-LCP vs. HPD-LCP P=1.30×10^−3^).

Given the large phylogenetic distances observed among co-active genomes (**Fig. 3a**) and communities (**Extended Data Fig. 7**), we sought to delineate distinct community types of co-active genomes in a non-supervised fashion (see methods). Using this approach, we distinguished four types of co-active genome communities: randomly-assembled communities, largely composed of genome communities with a high mean phylogenetic distance (PD) and a low metabolic cross-feeding potential (CP) score (HPD and LCP), which we used as a reference to define three other community types corresponding to two communities with a Low-PD (LPD) and High- or Low-CP (H/LCP), and a third community with High-PD (HPD) and High-CP (HCP) (**Fig. 4b**). These four co-active genome community types displayed distinct taxonomic compositions, with LPD-HCP communities mainly composed of Gamma- and Alphaproteobacteria, while HPD-HCP were more diverse including genomes from classes Nitrososphaeria, Marinisomatia, Dehalococcoidia, Alphaproteobacteria, and Acidimicrobiia (**Extended Data Fig. 9**). As anticipated, both HPD communities (orange and pink) were more dissimilar to respective LPD communities (blue and green) with regards to their encoded metabolism proxied by their functional Gini coefficient from KO genes occurrence profiles (**Fig. 4c**). Here, we hypothesised that these four community types displayed distinct signatures of metabolic exchanges and cross-feedings, which we analysed in details below.

### Key metabolic cross-feedings driving bacterioplankton community assembly

To further explore and identify molecular mechanisms driving these global patterns of predicted metabolic interactions, we analysed predicted metabolic exchanges within the four co-active genome community types delineated above. Both HPD-HCP and LPD-HCP communities were predicted to have a higher potential exchange in specific metabolites as revealed by a NMDS analysis of large metabolic categories (see methods) preferentially exchanged within each community type (**Fig. 5a**). Here, the first two dimensions of co-variation (Dim1 and Dim2) highlighted amino acids (AAs), B-vitamins, organo-sulfur compounds, aliphatic amines, n-alkanals, and aromatics as metabolic categories most preferentially exchanged within HPD-HCP and LPD-HCP co-active genomes community types (**Fig. 5b**). Despite large differences in mean PD within these communities, preferentially exchanged metabolic categories appeared to be conserved in HPD-HCP and LPD-HCP community types, suggesting these predicted metabolic exchanges are ancient and evolutionarily conserved^13^. This observation raises a key question regarding which evolutionary mechanisms can actually stabilize metabolic cross-feedings within natural microbial communities^56^. Although little is known about the coevolutionary consequences of cooperative cross-feeding, stable coevolution is expected to increase productivity in cross-feeding communities, which was corroborated by experimental evidence^57^. Zooming on large metabolic categories, we identified specific metabolites predicted to be preferentially exchanged within all four community types (**Fig. 5c**). When considering inorganic compounds for community metabolic modelling, most preferentially exchanged compounds among all community types were phosphate and iron cations (**Extended Data Fig. 10**), likely due to the essential uptake of these limiting nutrient and co-factors in the ocean^46^. Thus, in order to focus on actual biotic metabolic exchanges predicted, we did not consider inorganic compounds as previously done in other studies^25^.

**Figure 5:**
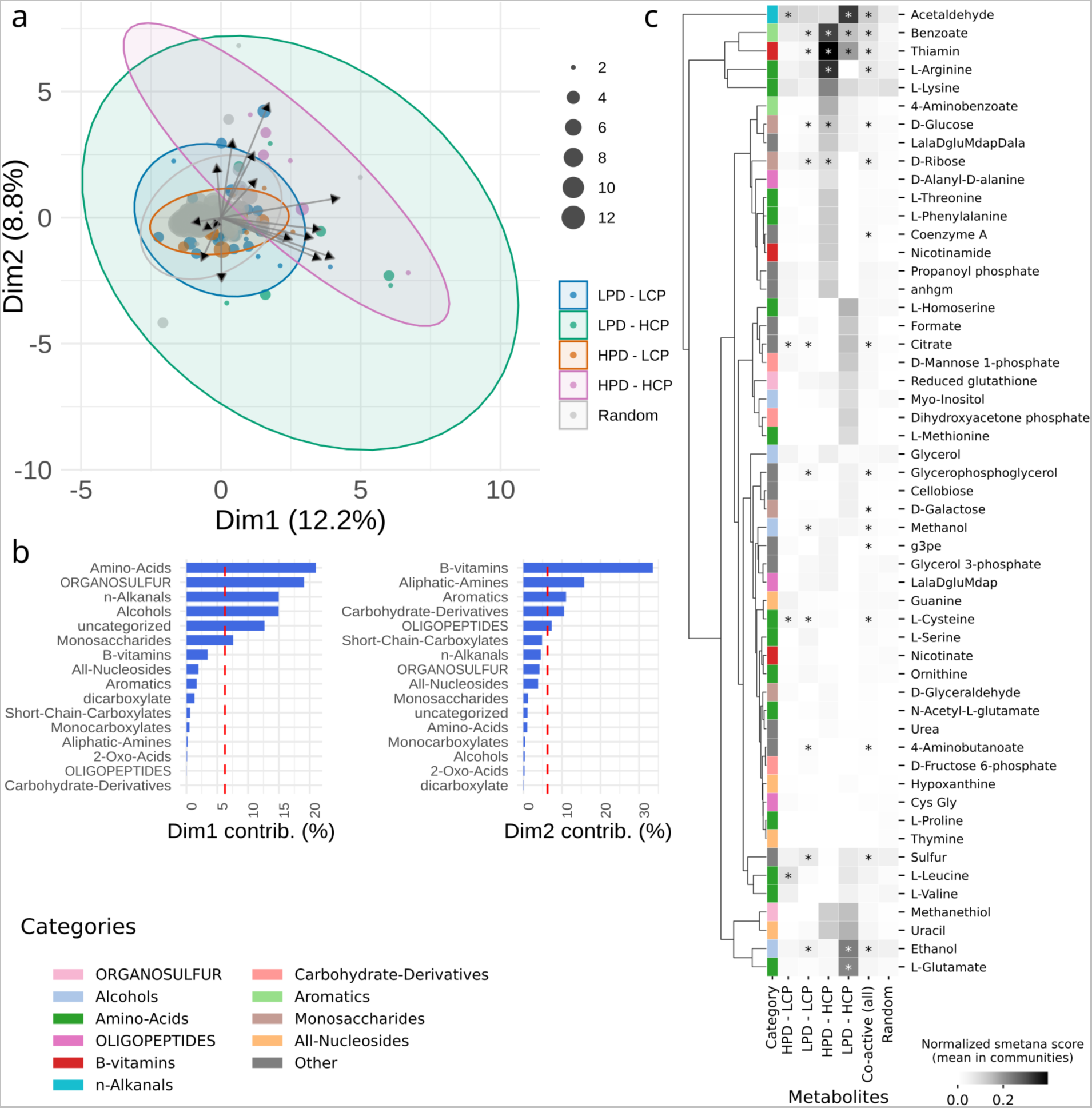
Community metabolic modelling predicts specific metabolic cross-feedings within co-active marine prokaryotic communities. **a,** A NMDS analysis revealed that HPD-HCP and LPD-HCP communities are predicted to have a higher potential exchange in specific metabolic categories. **b,** Overall, the higher potential for exchanges in HPD-HCP and LPD-HCP communities is driven by specific metabolic categories (NMDS Dim1 and Dim2), in particular amino acids, B vitamins, organo-sulfur compounds, and aliphatic amines. **c,** Within these large metabolic categories, specific metabolite exchanges are identified within each co-active genome community type. In particular, exchanges of acetaldehyde, benzoate, thiamin (vitamin B_1_), ethanol, and L-glutamate are predicted in LPD-HCP, while in HPD-HCP exchanges of benzoate, thiamin, L-arginine, as well as D-glucose and D-ribose are predicted.

Considering detailed predicted metabolic exchanges (using SMETANA sum scores) we identified compounds that were preferentially exchanged within each community type (**Fig. 5c and Supplementary Table 6**). In particular, acetaldehyde, benzoate, thiamine (vitamin B_1_), ethanol, and L-glutamate exchanges were enriched in LPD-HCP communities, while in HPD-HCP communities preferential exchanges of benzoate, thiamine, L-arginine, as well as D-glucose and D-ribose were predicted (**Supplementary Table 7**). The relative importance of predicted AA exchanges, and in particular biosynthetically costly AAs (e.g., methionine, lysine, leucine, arginine), likely reflects the key role of syntrophic interactions enabling cooperative growth in scarce environments^56^. Such division of metabolic labour for AAs can promote a growth advantage for cross-feeding species, as the fitness cost of overproducing AA has been experimentally shown to be less than the benefit of not having to produce them when they were provided by their partner^58^. Considering predicted L-glutamate exchanges, glutamic acids have been reported as potential auxophores (i.e., a compound that is required for growth by an auxotroph) in aquatic environments^59^. Notably, arginine and glutamate are linked in Cyanobacteria^60^ and plants^61^ through the metabolism of glutamate that involves the glutamate dehydrogenase for arginine synthesis, and which is an important network of nitrogen-metabolizing pathways for nitrogen assimilation. In marine microorganisms, nitrogen (N) cost minimization is an important adaptive strategy under global N limitation in the surface ocean, acting as a strong selective pressure on protein atomic composition^62^ and the structure of the genetic code^63^. Given that arginine plays an important role in the N cycle because it has the highest ratio of N to carbon among all AAs, the combined selective pressure at genomic level and for biosynthetic (N) cost minimization may explain the recurrent cross-feeding predictions of glutamate and arginine observed herein. Overall, these results support amino acid auxotrophy as a potential evolutionary optimizing strategy to reduce biosynthetic burden under nutrient (in particular N) limitation while promoting cooperative interactions^56,64^.

B-vitamins, which are essential micronutrients for marine plankton^65^, are predicted here to significantly structure bacterioplankton community activity, which supports the hypothesis that B-vitamin mediated metabolic interdependencies contribute to shaping natural microbial communities^66^. A recent environmental genomes survey in estuarine, marine, and freshwater environments has revealed that most naturally occurring bacterioplankton are B_1_ (thiamine) auxotrophs^67^. Vitamin interdependencies and auxotrophies, in particular for thiamine, have been recently predicted through a metagenomics-based association network in a soil microbial community, and confirmed in microcosm experiments^68^. Another comparative genomics assessment of vitamin B_12_ (cobalamin) dependence and biosynthetic potential in >40,000 bacterial genomes predicted that 86% of them require the cofactor, while only 37% encode a complete biosynthetic potential, the others being split into partial producers and salvagers^69^. In addition to thiamine, the joint importance in the metabolite exchanges of ornithine, glutamate and methionine, which are all products of enzymes dependent on vitamin B_12_^70^, confirms that access to vitamin B_12_ plays a significant role in structuring microbial community interactions. Furthermore, acetaldehydes are known intermediates supporting prokaryotic growth after breaking down substrates such as ethanolamine and propanediol using metabolic pathways involving vitamin B_12_-dependent enzymes^71^. Taken together, our results thus support the prevalent reliance of bacterioplankton on exogenous B_1_ and B_12_ precursors/products and on the bioavailability of micronutrients as important factors influencing bacterioplankton growth and community assembly.

Given the identification of amino acids, B vitamins and associated product exchanges as key metabolic mediators driving bacterioplankton community assemblies, we investigated their graph centrality within the co-activity network of bacterioplankton communities using the closeness centrality metric. The closeness centrality measures nodes centrality in a network by calculating the reciprocal of the sum of the length of the shortest paths between the node and all other nodes in the graph. The more central a node is, the closer it is to all other nodes. Overall, this revealed that genome donors, in particular for amino acids and B vitamins, displayed significantly higher closeness centrality than non-donor genomes (**Extended Data Fig. 11**). This observation supports the hypothesis that donor genomes influence community assembly via cross-feeding interactions through more central positions or hubs in the ecological network. Given that metabolic interdependencies predicted here are mainly observed among co-active genomes that are overall smaller in size (**Extended Data Fig. 5**), we also compared the genome sizes of donor vs. non-donor genomes, which revealed that non-donor genomes tended to be significantly smaller in size as compared to donor genomes (**Extended Data Fig. 11**). This observation actually supports the Black Queen Hypothesis (BQH)^72,73^, stating that species can gain a fitness advantage through genome streamlining, which is often observed (including herein) within marine bacterioplankton genomes^74^. Genome streamlining can reduce the nutrient requirements associated with the maintenance of more genetic material and limits energetically costly metabolic activities. Although our prediction results underline the key role of metabolic cross-feeding supporting positive interactions between microbes, many microorganisms in nature are prototrophic and are able to grow on simple substrates without the help of others^75^. Trade-off mechanisms such as resource allocation, design constraints, and information processing, can concomitantly shape microbial traits in the wild and lead to different biological adaptations leading to generalist or specialist lifestyles^76^. However, recent experimental work recently demonstrated that obligate cross-feeding can significantly expand the metabolic niche space of interacting bacterial populations^3^, thus potentially positively selecting cross-feeding bacterial populations.

The metabolic cross-feedings and interdependencies predicted here can be extremely useful to draw hypotheses for testing in the laboratory, for example through co-culture experiments. Focusing on one of the most abundant photosynthetic organisms on Earth, the marine cyanobacteria *Prochlorococcus sp.*, we further analysed predicted exchanges within a small community of six genomes (‘*coact-MHQ-014*’, see **Supplementary Table 6**) including one genome of *Prochlorococcus marinus*, three genomes of Pelagibacteraceae (two *Pelagibacter sp.* and one *MED-G40 sp.*), one genome of order Rhodospirillales (family UBA3470), and one genome of phylum Dadabacteria (*TMED58 sp.*). The community biogeography of this consortium revealed a globally distributed activity in both SRF and DCM, but restrained to mainly Westerlies (temperate) stations between 30° to 60° in absolute latitude (mean 33.8°N/27.4°S in SRF, mean 34.3°N/21.7°S in DCM) (**Extended Data Fig. 12**). Most robustly predicted exchanges within this community included the exchanges of several amino acids (L-arginine, L-homoserine, L-lysine, and L-phenylalanine), of vitamin B_1_ provided by a *Pelagibacter sp.* to two other genomes (*MED-G40 sp.* and family UBA3470), but also of D-ribose provided by the Rhodospirillales genome (family UBA3470) to *Prochlorococcus marinus*. The latter prediction provides a putative mechanism by which heterotrophic bacteria (such as from the order Rhodospirillales) can facilitate the growth of *Prochlorococcus marinus*^77^. While these metabolic exchanges remain predictions, they readily allow to formulate novel hypotheses to be further validated in the lab through co-culture experiments.

## Conclusion

In sum, these results underline the global-scale importance of trophic interactions influencing the co-activity, assembly, and resulting community structure of marine bacterioplankton communities^2^. Our computational predictions support in particular amino acids and B vitamin auxotrophies^29,67^ as important mechanisms driving bacterioplankton community assembly in the surface ocean. Given that these metabolic interdependencies are mainly observed among co-active genomes that are overall smaller in size, these results also support the Black Queen Hypothesis^72^ as an important mechanism shaping bacterioplankton community assembly in the global euphotic ocean. The integrated ecological and metabolic modelling framework developed herein has revealed the genomic underpinnings of predicted metabolic interdependencies shaping bacterioplankton community activity and assembly in the global surface ocean. It also revealed putative trophic metabolic interactions occurring among the most abundant bacterioplankton cells in the ocean (i.e., *Prochlorococcus* and *Pelagibacter*). Ultimately, these *in silico* predictions will have to be validated experimentally, through (high-throughput) co-culturing^78^. Finally, the computational framework developed here can readily be applied to the study of other microbiomes, in which mechanistic predictions of biotic interactions may also serve for generating novel hypotheses for co-culturing, with the goal to better capture the vast uncultivated microbial majority across microbial ecosystems. Overall, this framework integrating ecosystem-scale meta-omics information through ecological and metabolic modelling paves the way towards an improved functional and mechanistic understanding of microbial interactions driving ecosystem functions *in situ*.

## Methods

### A database of species-level marine prokaryotic genomes

A database of genomes from marine prokaryotes was assembled using several specialised databases as well as genomes reconstructed within specific studies. These databases included whole-genome sequences from marine prokaryote isolates (WGS), single-amplified genomes (SAGs), and metagenomic-assembled genomes (MAGs). The main database source for our genome collection was the Marine Metagenomic Portal^79^ through the use of the databases MarRef v4.0 (N=943, mostly high-quality WGS)^79^, MarDB v4.0 (N=12,963)^79^, and aquatic representative genomes from the ProGenomes database v1.0^32^ (N=566). This collection of well-documented genomes was complemented by 5,319 MAGs assembled from four distinct studies, namely: Parks et al. 2017^80^ (N=1,765; downloaded from EBI), Tully et al. 2017^81^/2018^20^ (N=2,597; downloaded from EBI), and Delmont et al. 2018^21^ (N=957; downloaded from FIGSHARE). The Parks et al. study contained genomes reconstructed from non-marine biomes. Thus, a selection of 1,765 genomes was extracted by searching for specific keywords: “tara, marine, sea, ocean, mediterranean” (case insensitive). Note that depending on their study of origin, included MAGs may have been reconstructed using different assembling and binning methods. Details about included genomes and their origins are reported in **Supplementary Table 1**. Overall, our marine genomes catalogue contained 19,791 highly redundant genomes (WGS, MAGs and SAGs). Genomes from this non-dereplicated catalogue were further filtered and quality-controlled before their inclusion in our study. We used CheckM v1.0.18^82^ to estimate the quality of the 19,791 genomes in our marine genomes catalogue (see <u>SnakeCheckM</u> in ecosysmic repository). Through the annotation and counting of single-copy marker genes (SCGs), CheckM estimates the level of completeness, contamination, and strain heterogeneity of individual genomes. We used those metrics to classify our genomes into three categories: high-quality (HQ) for ≥90% completeness ≤5% contamination (N=8,736), medium-to-high-quality (MHQ) for ≥75% completeness ≤10% contamination (N=4,547), and medium-quality (MQ) for ≥50% completeness ≤25% contamination (N=5,381). Genomes that did not meet at least the MQ threshold were tagged as low-quality (LQ) and discarded from the database (N=1,127). Quality estimates were used in the de-replication process that was performed using dRep v2.2.3^83^ (see <u>dReplication</u> in ecosysmic repository). dRep uses average nucleotide identity (ANI) and filters out redundant genomes *via* a 2-step clustering strategy: a fast coarse-grained clustering by MASH ANI (threshold used: 90% ANI over 60% of the genomes), followed by a slow fine-grained clustering through NUCMER ANI in clusters identified in the previous step only (threshold used: 95% ANI over 60% of the genomes). This process yielded 7,658 non-redundant species-level genomes with an average nucleotide identity below 95%, a threshold previously reported to delineate species level for prokaryotes^84^. These genomes were assigned taxonomic information using GTDB-TK v0.3.2^85^ (see <u>SnakeGTDBTk</u> in ecosysmic repository), which also allowed us to place our genomes within a phylogenetic tree using iTOL v5^86^. Since GTDB-Tk reconstructs two independent trees for Archaea and Bacteria, we linked them at the root using a distance of 0.122^87^, as recommended by the authors and tool maintainers (https://github.com/Ecogenomics/GTDBTk/issues/209).

### Functional annotations and reconstruction of genome-scale metabolic models

Coding DNA sequences (CDS) and proteins were inferred using Prodigal v2.6.3^88^ and annotated using eggnog-mapper v1.0 on the eggNOG v5.0^89^ orthology resource (see <u>GeneAnnotation</u> in ecosysmic repository). The sets of annotated genes were processed using CarveMe v1.5.1^53^ to reconstruct individual metabolic networks using the generic command “carve --output --universe -- nogapfill --fbc2 --verbose” (see <u>SnakeCarveMe</u> in ecosysmic repository). The template used for each top-down reconstruction (referred to as “universe” in the original CarveMe paper) was selected for each genome using the GTDB-Tk taxonomic assignments as either cyanobacteria, bacteria, or archaea. CarveMe was run without gap filling with the solver IBM CPLEX v12.10.

### Genomic scaling laws analysis

We used scaling laws as a framework to characterise the functional content of our genomic database (WGS, SAGs, MAGs). These genomic scaling laws were also used as a tool to properly identify enriched or depleted functional and metabolic potentials within specific groups of genomes (e.g., origin, co-active or not) by taking into account genome size and identifying enriched/depleted potential in proportion of the observed genome size. EggNOG provides 25 high-level categories and a KEGG Orthology (KO) equivalent for each Cluster of Orthologous Group (COG) annotation. The KO database also provides a 4-level hierarchy of (unnamed) functional categories. We were able to group our 23,224 KO identified in our catalogue into 54 high-level categories (level 2 in the hierarchy that presented for us the best compromise between specificity and tractability of the metabolic functions). For each high-level KO or COG category, we fitted a linear law on the log-transformed variables using the function scipy.stats.linregress v1.7.3 (parameter alternative=“greater”). Functional categories with a R^2^ below 0.3 were discarded, and the distribution of residuals were compared (in log-scale) using the Mann-Whitney U test using the function scipy.stats.mannwhitneyu v1.7.3 (parameter alternative=“two-sided”). P-values from all tests were corrected using Bonferroni and Benjamini-Hochberg multiple-testing corrections (see **Supplementary Table 2 and 4**) using the function stats.multitest.multipletests from the statsmodels Python package (v0.13.2).

### Functional Gini coefficient

In order to quantify how the functional potential of each community was shared between genomes, we used a proxy of the well-established Gini index. In Economics, the Gini index “measures the extent to which the distribution of income (or, in some cases, consumption expenditure) among individuals or households within an economy deviates from a perfectly equal distribution”. Inside each predicted co-active consortium, we defined a “functional capital” for each member as the sum of occurring KO that were present inside the genome, and computed the Gini index on this value. A Gini index of 0 can be interpreted as a perfect overlap between the functions of all members of the consortium, while a Gini index of 1 would be the extreme situation where a single member of the consortium displays all the detected KO functions. Intermediate values represent varying degree of metabolic evenness between the members of the community, a measure that we tried to use to separate niche overlap from potential metabolic complementarity.

### Meta-omics profiling and associated environmental contextual data

We leveraged metagenomics and metatranscriptomics data from samples of the *Tara* Oceans expeditions (2009–2013)^90^. We focused on samples from prokaryotic-enriched size fractions (0.2-1.6 μm and 0.22-3 μm) in the euphotic zone, including surface (SUR) and deep-chlorophyll maximum layer (DCM) samples. This yielded 107 samples across 64 stations for metagenomics data, 118 samples across 81 stations for metatranscriptomics data, and 71 samples across 45 stations for which we had both. Sequencing reads were previously quality-controlled using methods described in ^90^. We then mapped quality-controlled reads onto our 7,658 non-redundant marine prokaryotic genomes using Bowtie 2 v2.3.4.3^91^ (see <u>ReadMapping</u> in ecosysmic repository) using the command “bowtie2 -p --no-unal -x -1 -2 -S” with no extra parameter. Reads that successfully mapped were subsequently filtered using Samtools v1.9^92^ and pySAM v0.15.2 using MAPQ ≥ 20 and a nucleotide identity ≥ 95% to avoid non-specific mappings. The identity score ignores ambiguous bases (N) on the reference but takes gaps into account. The formula used is (NM - XN) / L with NM the edit distance; that is, the minimal number of one-nucleotide edits (substitutions, insertions and deletions) needed to transform the read string into the reference string, XN the number of ambiguous (N) bases in the reference, and L the length of the read. Overall, this ensured that the conserved reads were mapped to the target genome with a high-specificity. We estimated depth of coverage (i.e., vertical coverage) by dividing the total mapping of a genome by its size, and breadth of coverage (i.e., horizontal coverage) by dividing the number of mapped bases (at least one time) by the genome size (see <u>CoverageEstimation</u> in ecosysmic repository).

### Co-abundance and co-activity networks inference

Co-abundance and co-activity networks were reconstructed using *FlashWeave* (FW) v0.18.0^50^. FW relies on a local-to-global learning framework and infers direct associations by searching for conditional dependencies between features. Several heuristics are then applied to connect these local dependencies and infer a network. We defined the abundance of a genome in a sample by its overall metagenomic vertical coverage (also called depth) per 1M base pairs, while its activity was given by the ratio of its overall metatranscriptomic coverage depth per 1M base pairs over its abundance. Note that this can only be computed at stations and depths for which we have both metagenomic and metatranscriptomic signals. A given genome was defined as observed (i.e., present and/or active) within a sample when at least 30% of its genome was horizontally covered (also called breadth).

Overall, we were able to compute abundances for 107 samples, and activities for only 71 samples. To lower spurious correlations, abundance and activity data points for unobserved genomes were discarded and genomes with less than 10 observations across our samples were removed. This was done independently for abundance (N=1,232 genomes observed in at least 10/71 samples) and activity (N=902 genomes observed in at least 10/71 samples). Finally, the inherent compositional nature of the sequencing datasets was taken into account using centred log-ratio (CLR) transformation and the adaptive pseudo-count implemented in *FlashWeave*. Both abundance and activity matrices were used as input to *FlashWeave* using parameters “normalize=true, “n_obs_min=10, max_k=3, heterogenous=true” (see the FlashWeave documentation for more information about these parameters). Genome graph centralities were computed with the *networkx* python library v3.1 using the *closeness_centrality* function on the co-activity community networks for which metabolic exchanges were predicted using SMETANA (see below).

### Community metabolic modelling and cross-feeding interaction predictions

We identified co-active genome communities in the reconstructed co-activity network using the Markov clustering algorithm^93^ (MCL) through the use of *run_mcl* function with an inflation parameter of 1.5 available in Python *markov_clustering* library V.0.0.2. We also generated randomly-assembled communities by randomly sampling genomes from the pool of genomes used for network reconstruction (genomes occurring at least 10 times within the considered samples). These communities were quality-filtered for MHQ+HQ genomes and analysed using SMETANA 1.2.0^55^ to predict putative metabolic cross-feeding interactions (see <u>SnakeMETANA</u> in ecosysmic repository). SMETANA does not use any biological objective functions and is formulated as a mixed linear integer problem (MILP) that enumerates the set of essential metabolic exchanges within a community with non-zero growth of all community species subject to mass balance constraints. We limited the community metabolic analyses to MHQ+HQ genomes in order to lower the risk of predicting spurious interactions in communities of lower-quality genomes and metabolic models. SMETANA was run in both global and detailed modes with the solver IBM CPLEX v12.10, using in each mode the default media provided by the package (which is a complete media for global analysis, and a community-specific minimal media for detailed analysis). A set of inorganic compounds were excluded from the analysis as explicitly recommended by one of the package author (https://github.com/cdanielmachado/smetana/issues/20#issuecomment-827389107). Other parameters used were “--flavor bigg --solver CPLEX --molweight”.

The “community smetana score” reported in the main text is obtained by summing all smetana scores predicted for a given community. In order to compare communities of different sizes, this score was normalised by dividing the “smetana score” by the total number of potential genome-genome interactions, i.e. N x (N-1) / 2 (with N the size of the community). We referred to this new score in the main text as “normalised smetana score”.

In order to classify the different metabolites in the SMETANA database into metabolite categories (e.g., amino acids, carboxylates), we first mapped the metabolite identifiers to the MetaNetX database (available at: https://www.metanetx.org/cgi-bin/mnxget/mnxref/chem_xref.tsv). From this mapping, we extracted MetaCyc identifiers to subsequently obtain their ontologies (available at: https://metacyc.org/groups/export?id=biocyc14-14708-3818508891&tsv-type=FRAMES).

## Statistical analyses

All statistical tests and analyses were performed using *scipy.stats* Python module v1.7.3. All figures were generated using Python v3.7.12 and R v4.2.2. We used statannotations v0.4.4 (https://github.com/trevismd/statannotations) to append statistical significance to all boxplots. Stars are used to define significance level as follow: **** for P ≤ 10^−4^, *** for 10^−4^ < P ≤ 10^−3^, ** for 10^−3^ < P ≤ 10^−2^, * for 10^−2^ < P ≤ 5×10^−2^, and finally ns for P > 5×10^−2^. All data analysis sub-packages were installed in the same environment using Conda v22.11.1, the versions of which are detailed in the yaml file located in each repository cited above.

## Data availability

All data associated with this study are available in the main text, the supplementary materials, and at zenodo: https://zenodo.org/record/7853699#.ZEQ8ahVBx0Q.

## Code availability

All code repositories cited below are available within https://gitlab.univ-nantes.fr/ecosysmic.

## Acknowledgements

Tara Oceans (which includes both the Tara Oceans and Tara Oceans Polar Circle expeditions) would not exist without the leadership of the Tara Ocean Foundation and the continuous support of 23 institutes (http://oceans.taraexpeditions.org). We wish to thank the commitment of the following sponsors: CNRS (in particular Groupement de Recherche GDR3280 and the Research Federation for the study of Global Ocean Systems Ecology and Evolution, FR2022/Tara Oceans-GOSEE), European Molecular Biology Laboratory (EMBL), Genoscope/CEA, The French Ministry of Research, and the French Government ‘Investissements d’Avenir’ programmes OCEANOMICS (ANR-11-BTBR-0008), FRANCE GENOMIQUE (ANR-10-INBS-09-08), the CNRS MITI through the interdisciplinary program Modélisation du Vivant (GOBITMAP grant to SC), the RFI ATLANSTIC2020 (ECOSYSMIC grant to SC), and the H2020 project AtlantECO (award number 862923). NG and ED were supported by the RFI ATLANSTIC2020 (ECOSYSMIC and PROBIOSTIC grants). We also thank the support and commitment of Agnès b. and Etienne Bourgois, the Prince Albert II de Monaco Foundation, the Veolia Foundation, Region Bretagne, Lorient Agglomeration, Serge Ferrari, World Courier, and KAUST. The global sampling effort was enabled by countless scientists and crew who sampled aboard the Tara from 2009-2013, and we thank MERCATOR-CORIOLIS and ACRI-ST for providing daily satellite data during the expedition. We are also grateful to the countries who graciously granted sampling permissions. Computational support was provided by the bioinformatics core facility of Nantes (BiRD - Biogenouest), Nantes Université, France. The authors declare that all data reported herein are fully and freely available from the date of publication, with no restrictions, and that all of the analyses, publications, and ownership of data are free from legal entanglement or restriction by the various nations whose waters the Tara Oceans expeditions sampled in. This article is contribution number XXX of Tara Oceans.

## Author information

These authors contributed equally: Nils Giordano and Marinna Gaudin.

## Contributions

S.C. designed the research. N.G., M.G., and S.C. analysed the data, performed bioinformatic analyses, analysed and interpreted the results. C.T. and E.D. analysed the data and performed bioinformatic analyses. N.G. and S.C. wrote the paper with inputs from all other authors.

## Extended Data

**Extended Data Figure 1:**
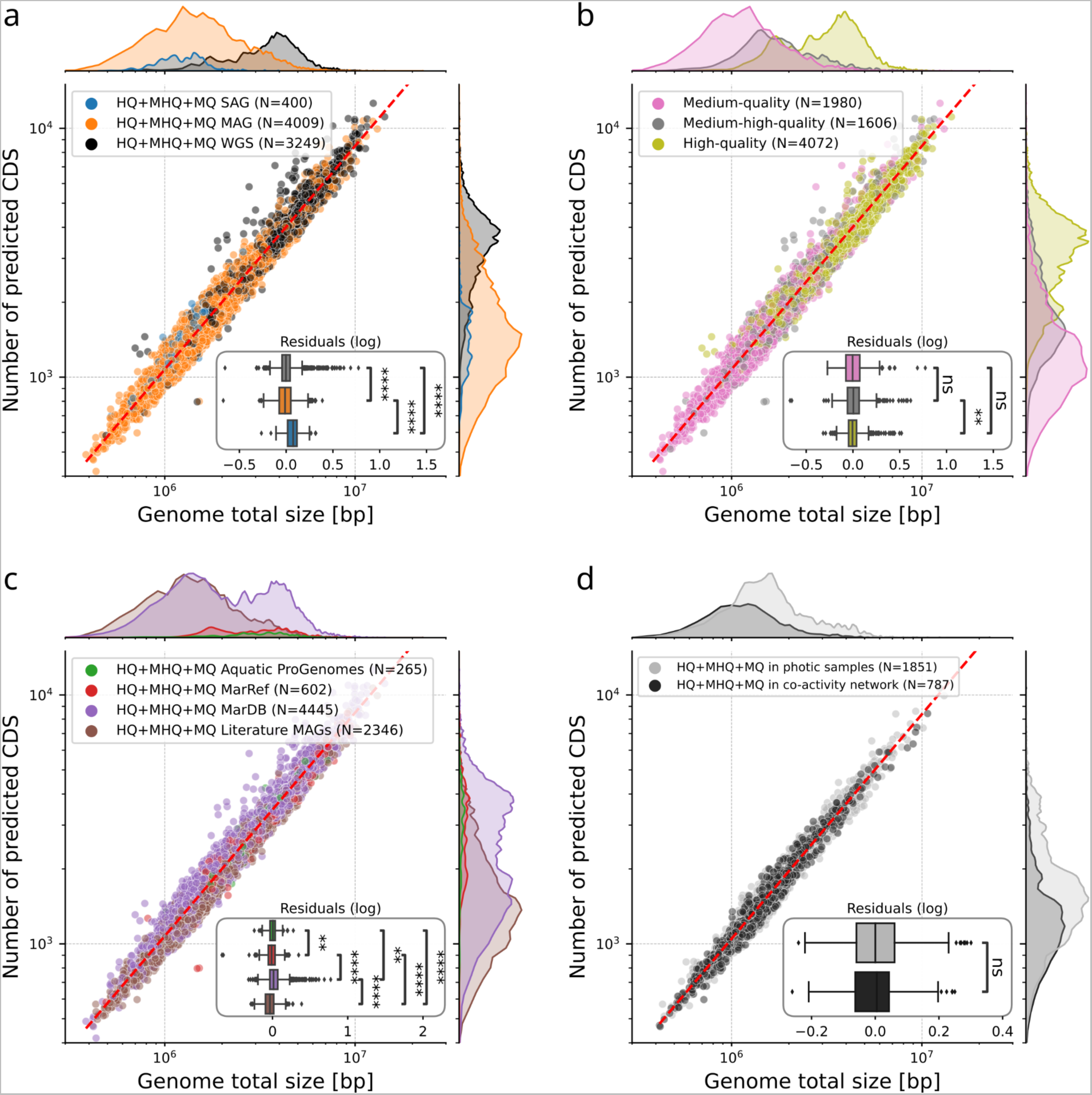
Genomics scaling laws for isolates and uncultivated prokaryotic genomes reconstructed from marine metagenomes. Comparison of genome size and number of predicted CDS for Medium Quality (MQ, completeness ≥ 50% and contamination ≤ 25%) dereplicated (95% ANI) genomes. We tested for significant deviations from the common scaling law by **a)** genome type (WGS, SAG, MAG), **b)** genome quality, **c)** source of genome, and **d)** presence in co-activity network (Mann–Whitney U on residuals with Bonferroni correction, best fit parameters and p-values are described in Supplementary Table 2).

**Extended Data Figure 2:**
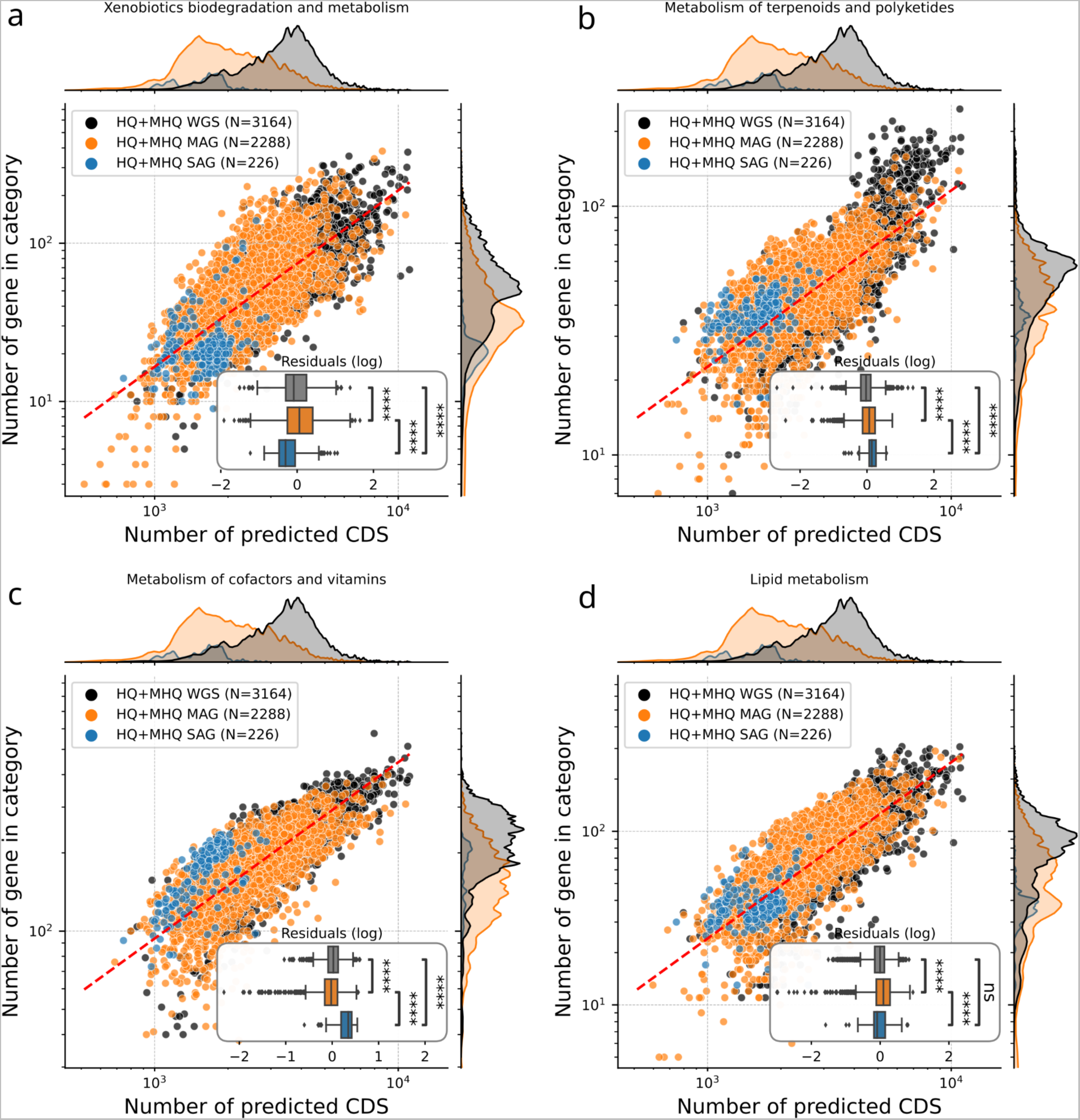
Scaling laws in the functional content of genomes for isolates and uncultivated prokaryotic genomes reconstructed from marine metagenomes. Abundance of annotated genes (KEGG database) coding for the metabolism of **a)** xenobiotics biodegradation, **b)** terpenoids and polyketides, **c)** cofactors and vitamins, and **d)** lipids, as a function of the number of CDS for Medium-High Quality and High Quality (MHQ+HQ, completeness ≥ 75% and contamination ≤ 10%) dereplicated (95% ANI) genomes. We tested for significant deviations from the common scaling law by genome type (Mann–Whitney U on residuals with Bonferroni correction, best fit parameters and p-values are described in Supplementary Table 2).

**Extended Data Figure 3:**
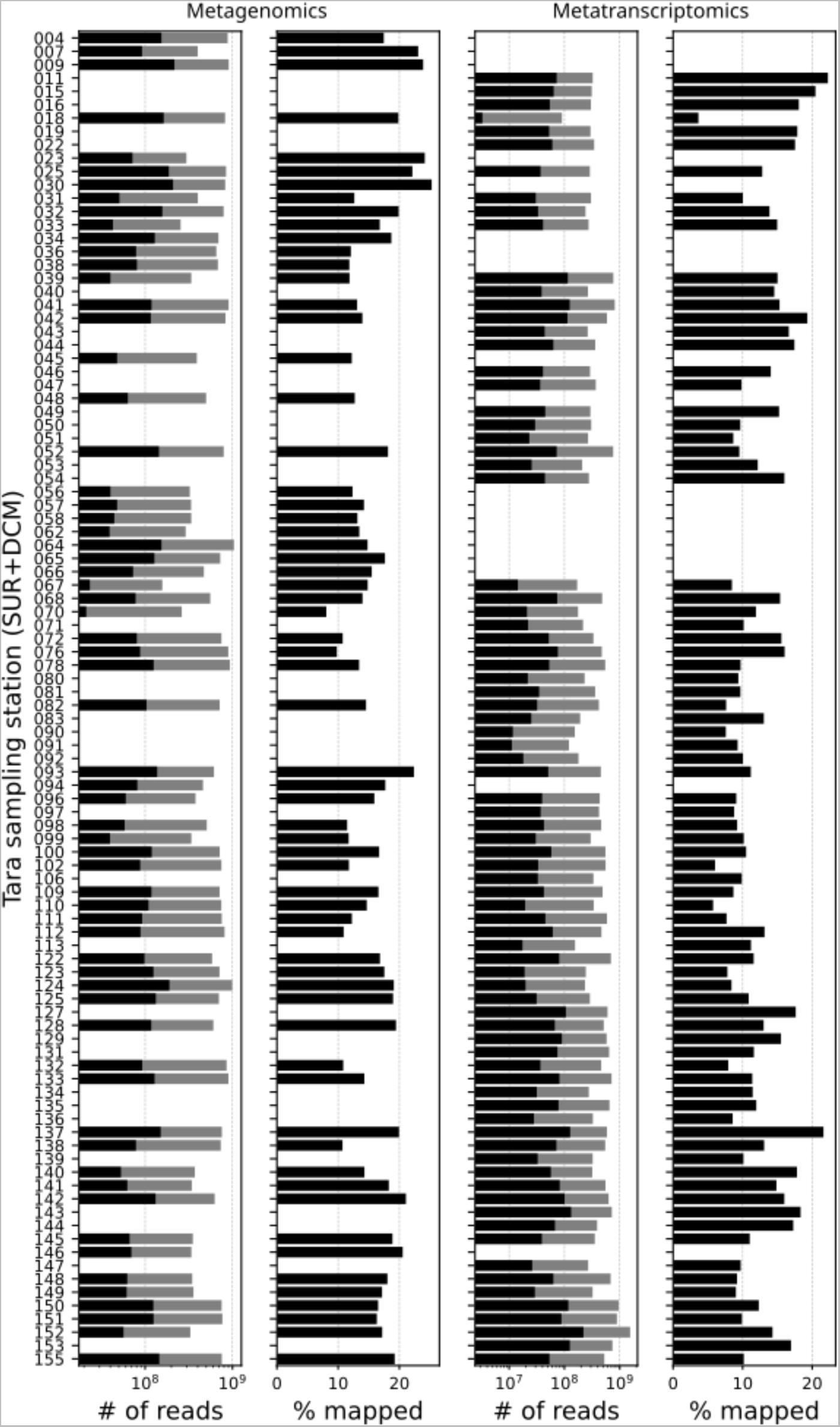
Mapped and Total reads for Metagenomics and Metatranscriptomics across *Tara* Oceans samples. Mapping results on the dRep95 catalogue for the metagenomics and metatranscriptomics euphotic samples from *Tara* Oceans expeditions (2009–2013). Black and grey bars are the number of mapped and total reads, respectively. Average mapping rates were 16.0% for metagenomes and 12.3% for metatranscriptomes. We used samples with both metagenomics and metatranscriptomics available to compute genome-wide co-activity.

**Extended Data Figure 4:**
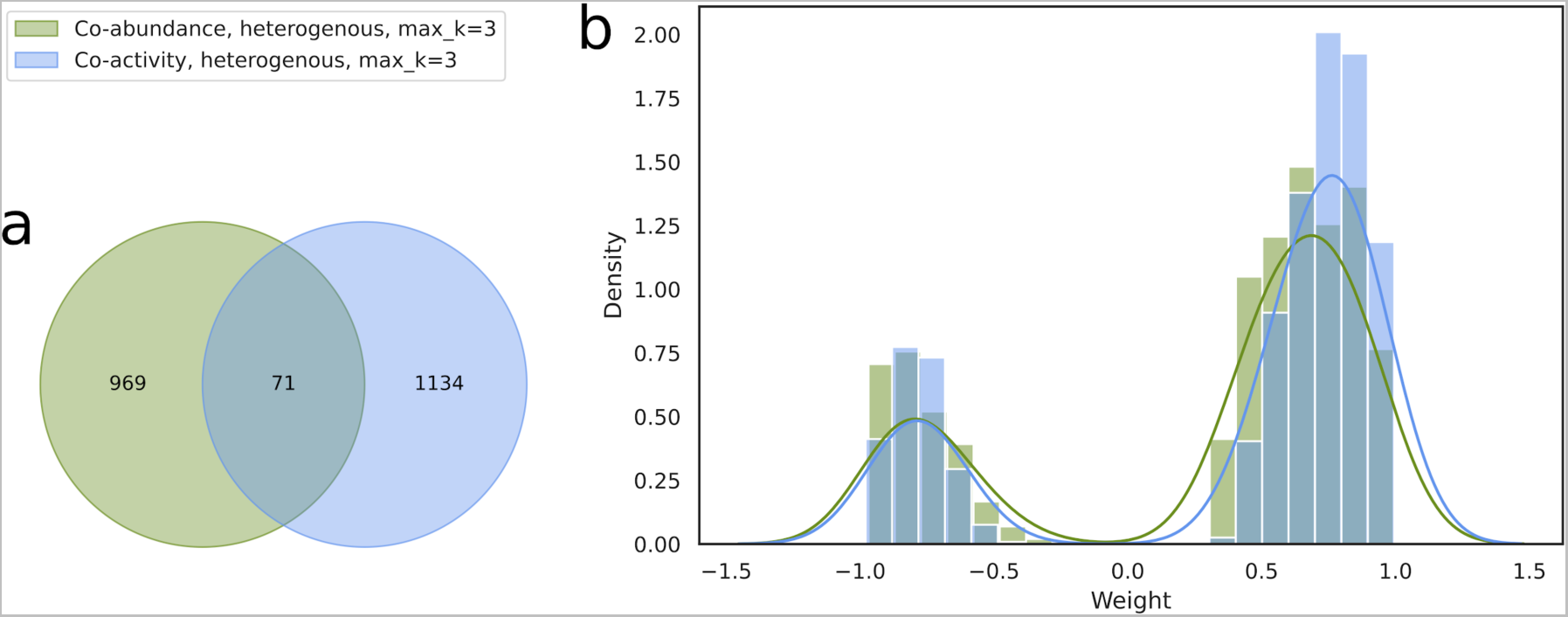
Comparison of genome-resolved co-abundance and co-activity networks. **a,** Venn diagram representing the number of shared and unique edges in the global genome-resolved co-abundance and co-activity networks. Only 71 associations were common to both networks, while 1,134 associations are specific to the co-activity network and 969 to the co-abundance network. **b,** Distributions of network weights (inferred by FlashWeave) in both networks. The co-activity network displayed significantly higher weights for positive associations as compared to the co-abundance network (Mann-Whitney U test, P < 0.001).

**Extended Data Figure 5:**
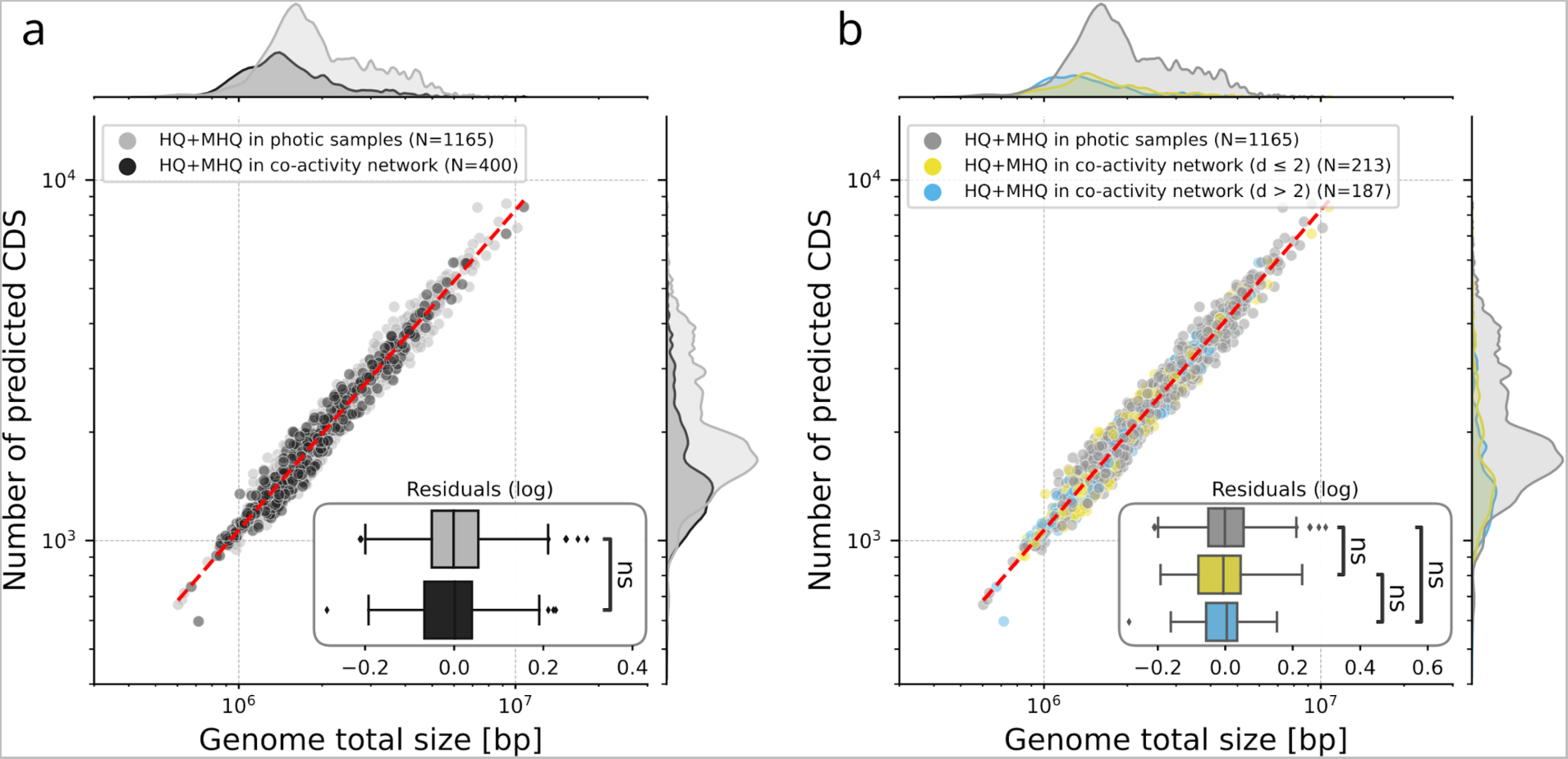
Genomic scaling laws for active and co-active genomes. Comparison of genome size and number of predicted CDS for Medium-High Quality and High Quality (MHQ+HQ, completeness ≥ 75% and contamination ≤ 10%) dereplicated (95% ANI) genomes. We tested for significant deviations from the represented log-log linear law by **a)** presence in the co-activity network, and **b)** below-median or above-median connectivity degree in the co-activity network (Mann–Whitney U on residuals with Bonferroni correction, best fit parameters and p-values are described in Supplementary Table 2). Genomes in photic samples are genomes that were detected active in at least one sample (see Methods). Genomes in the co-activity network are significantly smaller both in size and number of CDS (Mann–Whitney U test, p-value = 1.84 x 10^−45^ and 3.22 x 10^−46^ respectively).

**Extended Data Figure 6:**
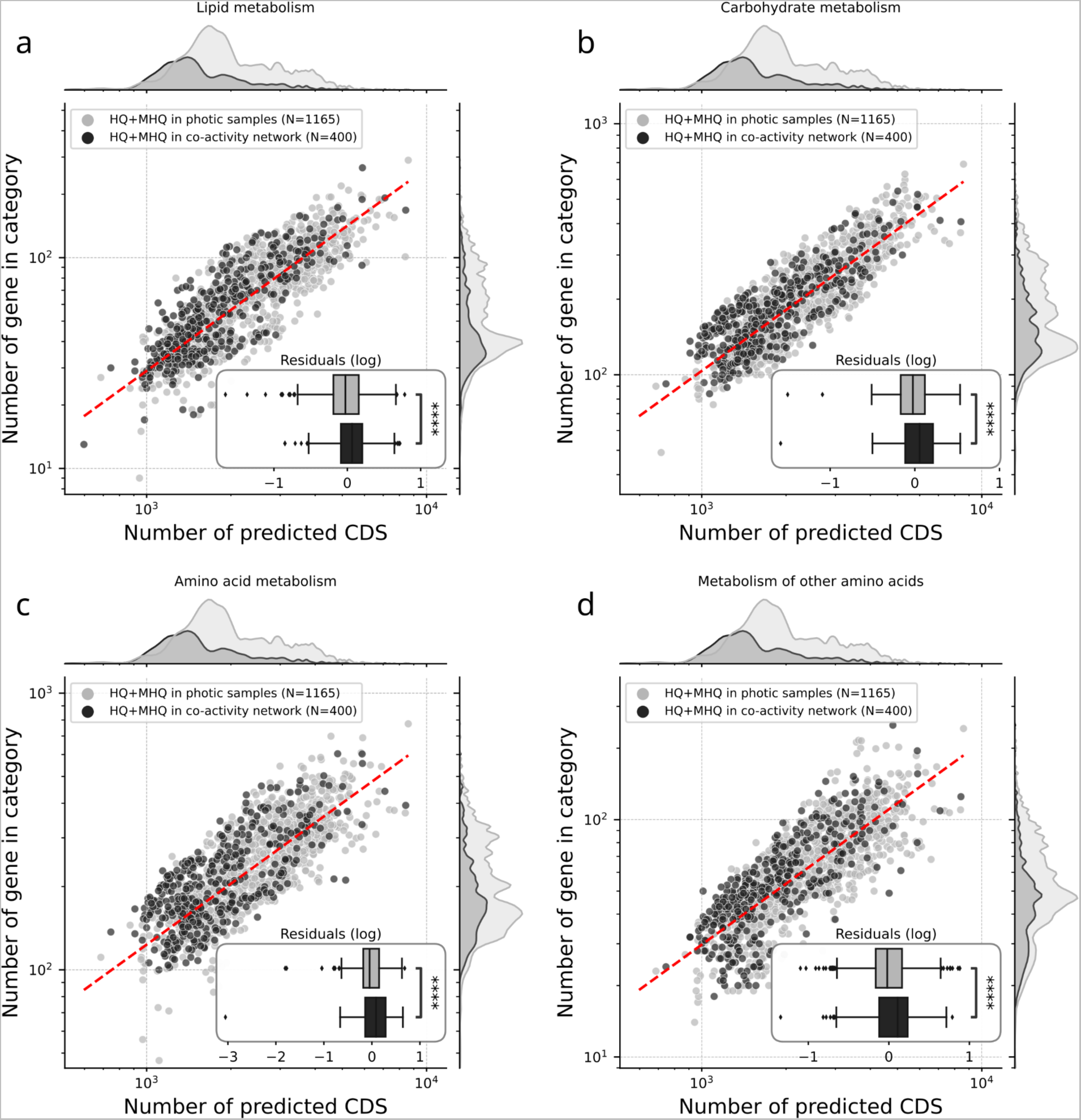
Scaling laws in the functional content of genomes for active and co-active genomes. Abundance of annotated genes (KEGG database) coding for the metabolism of **a)** lipids, **b)** carbohydrates, **c)** amino acids, and **d)** other amino acids, as a function of the number of CDS for Medium-High Quality and High Quality (MHQ+HQ, completeness ≥ 75% and contamination ≤ 10%) dereplicated (95% ANI) genomes. We tested for significant deviations from the common scaling law by category of genome (Mann–Whitney U on residuals with Bonferroni correction, best fit parameters and p-values are described in Supplementary Table 5).

**Extended Data Figure 7:**
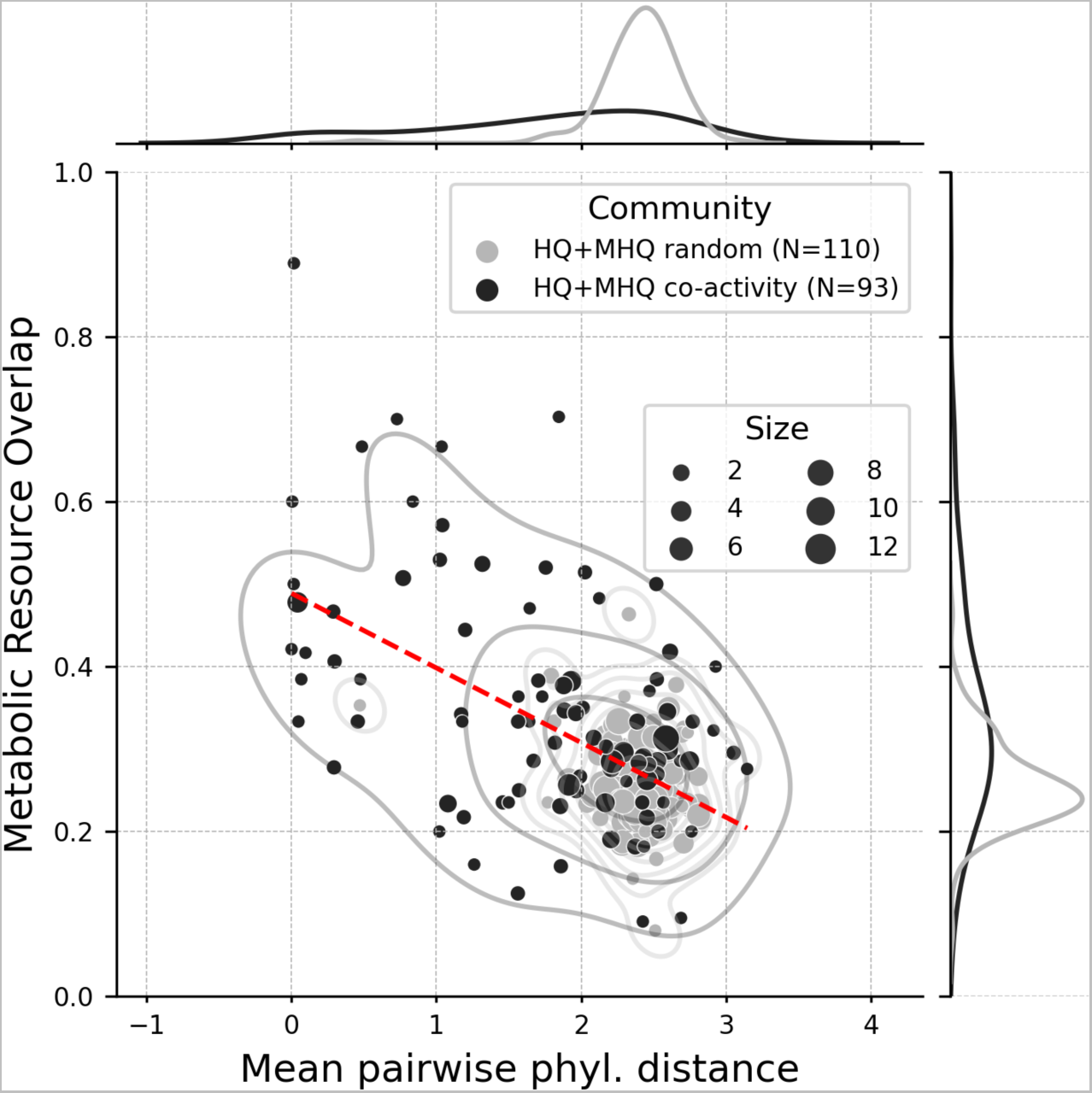
Metabolic resource overlap as a function of phylogenetic distance within communities of co-active genomes. Comparison of Metabolic Resource Overlap (SMETANA global score) and Mean Pairwise Phylogenetic Distance for co-active and randomly-assembled genome communities (see methods). Dashed-red line is the best linear fit and shows a significant negative relationship (slope=-0.091; intercept=0.49; r^2^=0.31; p-value=4.17 x 10^−18^).

**Extended Data Figure 8:**
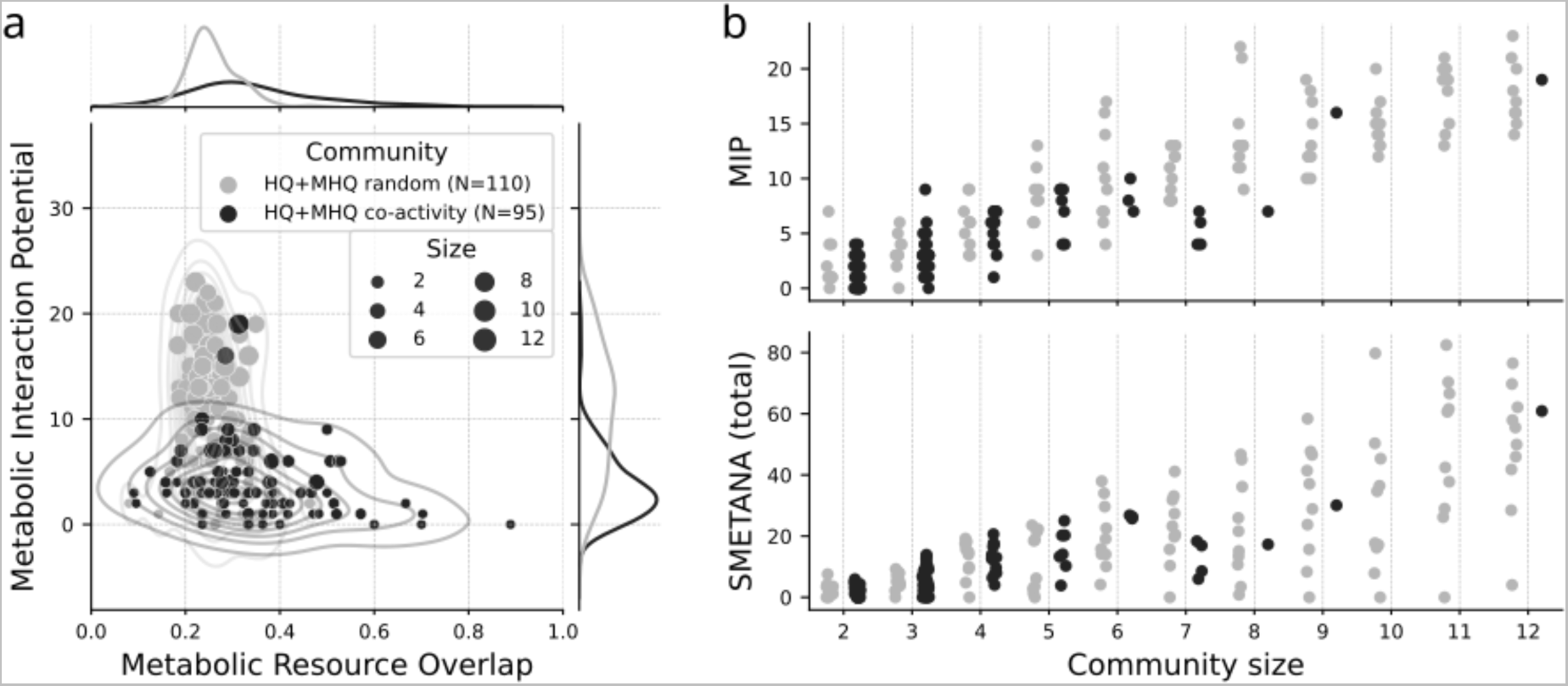
Community-wide metabolic modelling within marine prokaryotic communities. **a,** Comparison between Metabolic Interaction Potential (MIP) and Metabolic Resource Overlap (MRO) for co-active and randomly-assembled genome communities. A lower MIP score and a higher MRO score was observed for co-active genome communities as compared with randomly-assembled genome communities. **b,** Effect of community size on MIP and SMETANA scores for co-active and random communities. Both scores were significantly driven by community size (MIP R^2^=0.82, p-value=1.03×10^−77^; SMETANA R^2^=0.59, p-value=7.28×10^−41^).

**Extended Data Figure 9:**
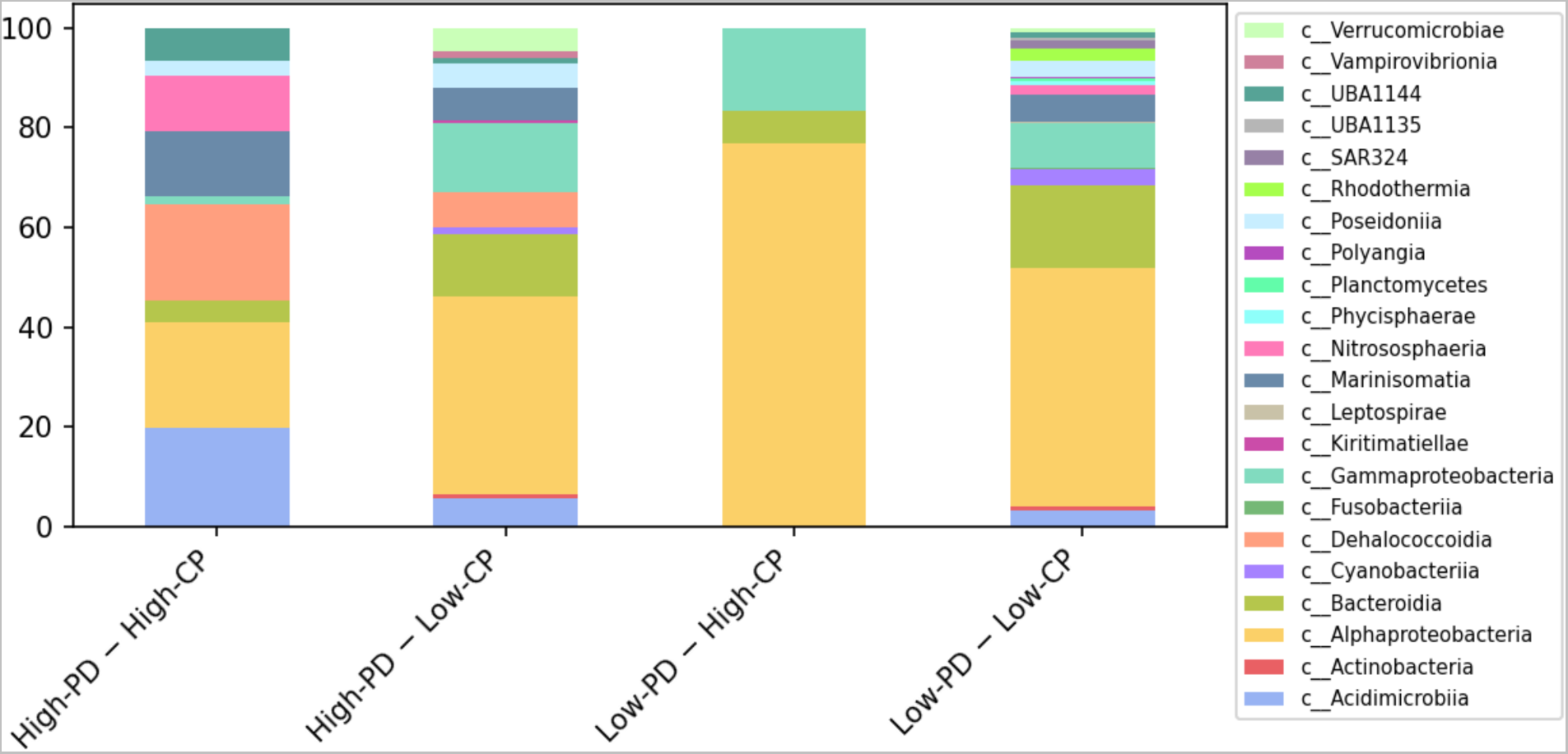
Taxonomic composition of co-active genome community types at the organismal class level. The taxonomic composition of the four co-active genome community types is presented as relative proportion at the class-level. The four co-active genome community types displayed distinct taxonomic compositions, with LPD-HCP communities mainly composed of Gamma- and Alphaproteobacteria, while HPD-HCP were more diverse including genomes from classes Nitrososphaeria, Marinisomatia, Dehalococcoidia, Alphaproteobacteria, and Acidimicrobiia.

**Extended Data Figure 10:**
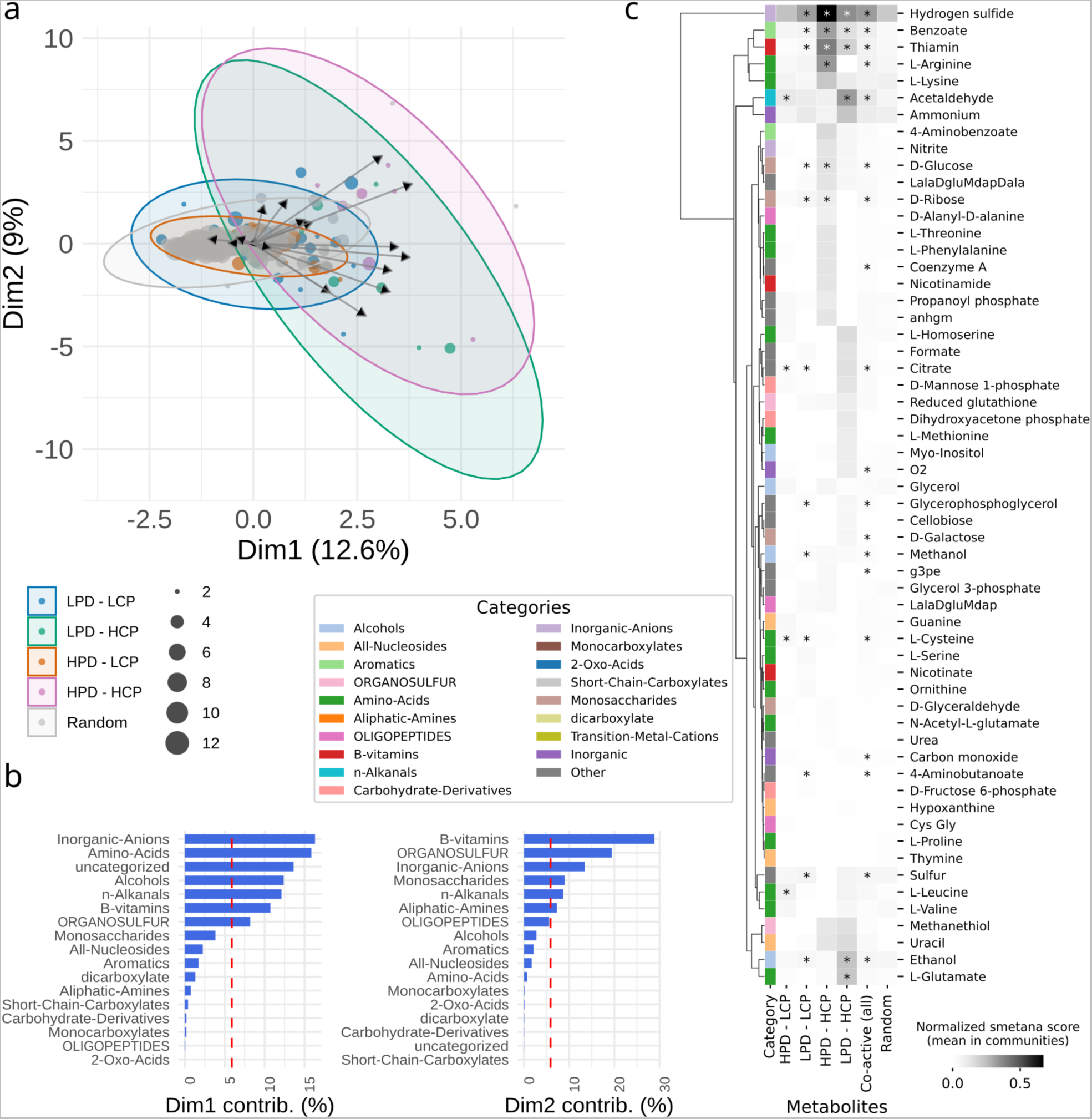
Detailed community metabolic modeling without considering inorganic compounds. **a,** NMDS analysis of the co-active genome communities in the space of putative metabolic exchanges (normalized smetana score) for each community type (defined in Fig. 4a). **b,** Contribution of high-level categories of metabolic compounds to the first two dimensions of the NMDS. **c,** Mean normalized smetana score for each high-level category of metabolic compounds in each community type. Stars denote a significant difference between categories (Mann-Whitney U, Benjamini-Hochberg correction, corrected p-value ≤ 0.05, all test results are available in Supplementary Table 7).

**Extended Data Figure 11.**
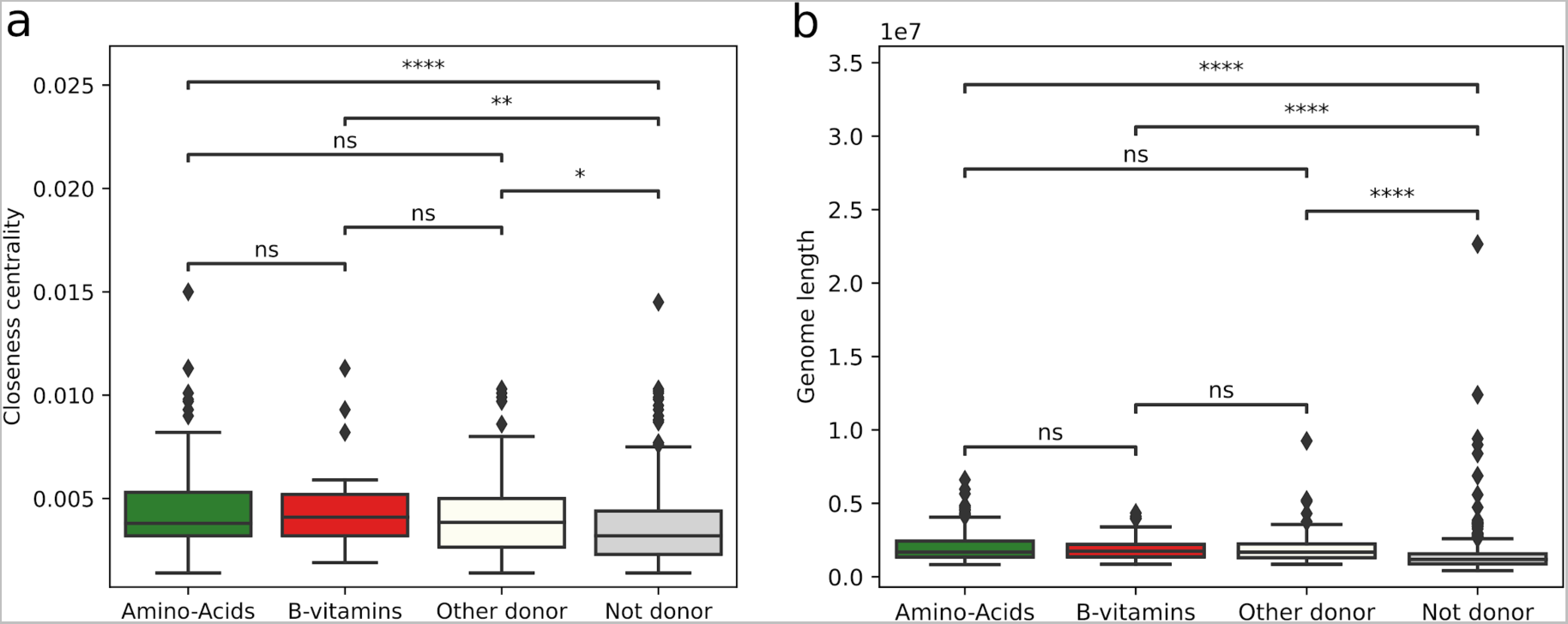
Co-active network closeness centrality and genome size for amino acids donors, B vitamins donors, other compounds donors, and non-donors. **a,** Closeness centrality estimates how fast the flow of information would be through a given node to other nodes. All categories of donors had a significantly higher closeness centrality index as compared to non-donors in the co-activity network (Mann-Whitney U test, Benjamini-Hochberg correction). **b,** Similarly, all categories of donors had significantly higher genome size as compared to non-donors (Mann-Whitney U test, Benjamini-Hochberg correction).

**Extended Data Figure 12.**
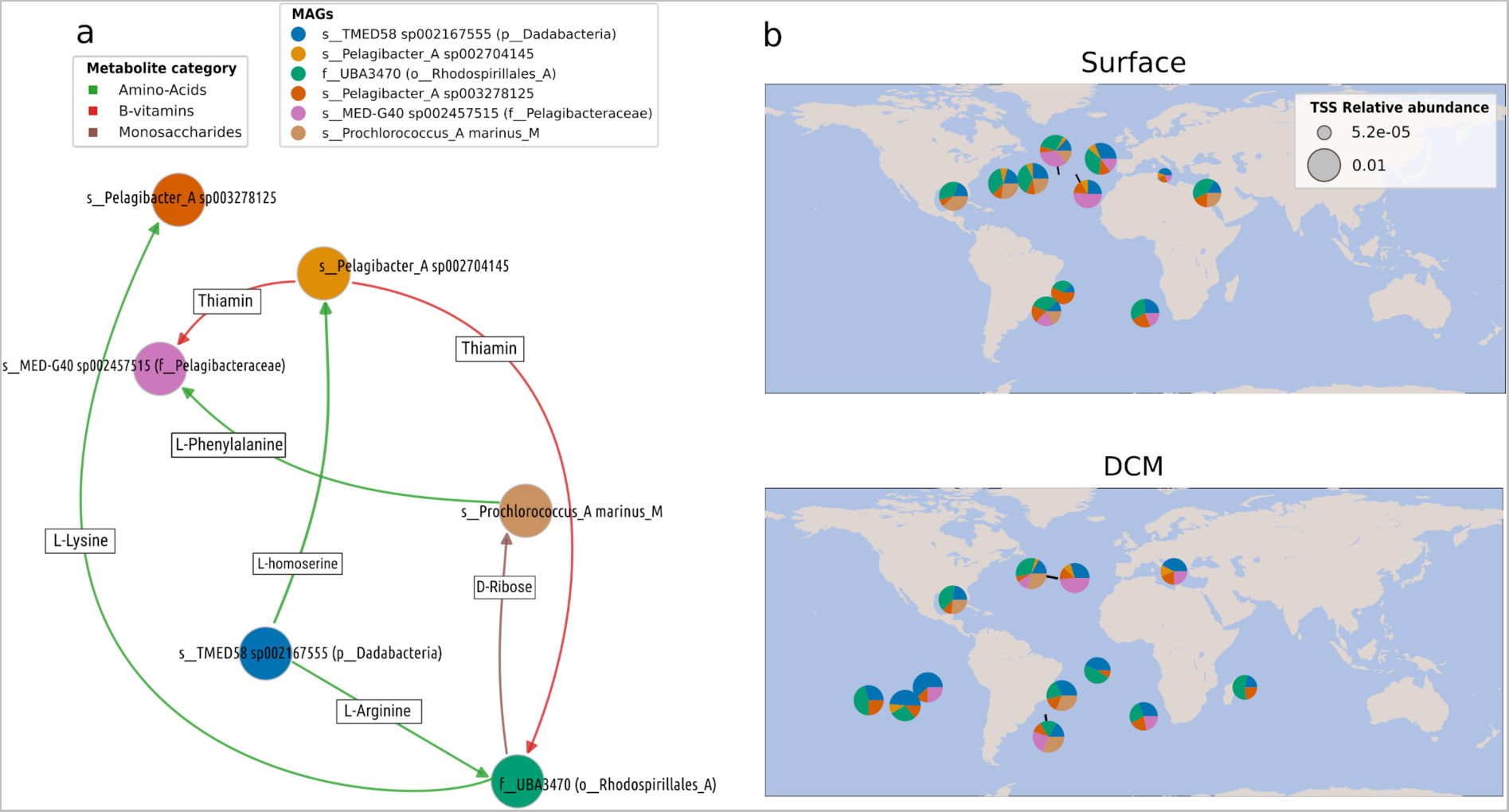
Zooming on a specific co-active genome community including a *Prochlorococcus marinus* genome: Predicted metabolic exchanges and biogeography. **a,** Graph representing predicted metabolic exchanges (SMETANA score >= 0.5) between genomes of community ‘*coact-MHQ-014*’. This community included one genome of *Prochlorococcus marinus* (brown), three genomes of Pelagibacteraceae (two Pelagibacter sp. and one MED-G40 sp.; red, gold and pink), one genome of order Rhodospirillales (family UBA3470; green), and one genome of phylum Dadabacteria (TMED58 sp.; blue). Exchanges of several amino acids, B1 vitamin, and D-Ribose were predicted between these genomes. **b,** Biogeography of the respective community ‘*coact-MHQ-014*’ and corresponding genome relative abundances at SRF and DCM *Tara* Oceans stations. The community was considered active if there were at least two genomes and one Pelagibacter detected at each station. The biogeography of this community revealed a globally distributed activity in both SRF and DCM, but restrained to mainly Westerlies (temperate) stations between 30°-60°N/S latitude (mean 33.8°N/27.4°S in SRF, mean 34.3°N/21.7°S in DCM).

## Supplementary information

Supplementary Tables 1–7

## References

1 Arrigo, K. R. Marine microorganisms and global nutrient cycles. Nature 437, 349–355, doi:10.1038/nature04159 (2005).

2 Gralka, M., Szabo, R., Stocker, R. & Cordero, O. X. Trophic Interactions and the Drivers of Microbial Community Assembly. Curr Biol 30, R1176–R1188, doi:10.1016/j.cub.2020.08.007 (2020).

3 Ona, L. et al. Obligate cross-feeding expands the metabolic niche of bacteria. Nat Ecol Evol 5, 1224–1232, doi:10.1038/s41559-021-01505-0 (2021).

4 Follett, C. L. et al. Trophic interactions with heterotrophic bacteria limit the range of Prochlorococcus. Proc Natl Acad Sci U S A 119, doi:10.1073/pnas.2110993118 (2022).

5 San Roman, M. & Wagner, A. Diversity begets diversity during community assembly until ecological limits impose a diversity ceiling. Mol Ecol 30, 5874–5887, doi:10.1111/mec.16161 (2021).

6 Evans, R. et al. Eco-evolutionary Dynamics Set the Tempo and Trajectory of Metabolic Evolution in Multispecies Communities. Curr Biol 30, 4984–4988 e4984, doi:10.1016/j.cub.2020.09.028 (2020).

7 Hug, L. A. et al. A new view of the tree of life. Nat Microbiol 1, 16048, doi:10.1038/nmicrobiol.2016.48 (2016).

8 Fritts, R. K., McCully, A. L. & McKinlay, J. B. Extracellular Metabolism Sets the Table for Microbial Cross-Feeding. Microbiol Mol Biol Rev 85, doi:10.1128/MMBR.00135-20 (2021).

9 Zengler, K. & Zaramela, L. S. The social network of microorganisms - how auxotrophies shape complex communities. Nat Rev Microbiol 16, 383–390, doi:10.1038/s41579-018-0004-5 (2018).

10 Faust, K. & Raes, J. Microbial interactions: from networks to models. Nat Rev Microbiol 10, 538–550, doi:10.1038/nrmicro2832 (2012).

11 Chaffron, S. et al. Environmental vulnerability of the global ocean epipelagic plankton community interactome. Sci Adv 7, doi:10.1126/sciadv.abg1921 (2021).

12 Blanchet, F. G., Cazelles, K. & Gravel, D. Co-occurrence is not evidence of ecological interactions. Ecology Letters 23, 1050–1063, doi:10.1111/ele.13525 (2020).

13 van den Berg, N. I., et al. Ecological modelling approaches for predicting emergent properties in microbial communities. Nat Ecol Evol 6, 855–865, doi:10.1038/s41559-022-01746-7 (2022).

14 Estrela, S. et al. Functional attractors in microbial community assembly. Cell Syst 13, 29–42 e27, doi:10.1016/j.cels.2021.09.011 (2022).

15 Pontrelli, S. et al. Metabolic cross-feeding structures the assembly of polysaccharide degrading communities. Sci Adv 8, eabk3076, doi:10.1126/sciadv.abk3076 (2022).

16 Sunagawa, S. et al. Tara Oceans: towards global ocean ecosystems biology. Nat Rev Microbiol 18, 428–445, doi:10.1038/s41579-020-0364-5 (2020).

17 Acinas, S. G. et al. Deep ocean metagenomes provide insight into the metabolic architecture of bathypelagic microbial communities. Communications Biology 4, 604, doi:10.1038/s42003-021-02112-2 (2021).

18 Larkin, A. A. et al. High spatial resolution global ocean metagenomes from Bio-GO-SHIP repeat hydrography transects. Sci Data 8, 107, doi:10.1038/s41597-021-00889-9 (2021).

19 Biller, S. J. et al. Marine microbial metagenomes sampled across space and time. Sci Data 5, 180176, doi:10.1038/sdata.2018.176 (2018).

20 Tully, B. J., Graham, E. D. & Heidelberg, J. F. The reconstruction of 2,631 draft metagenome-assembled genomes from the global oceans. Sci Data 5, 170203, doi:10.1038/sdata.2017.203 (2018).

21 Delmont, T. O. et al. Nitrogen-fixing populations of Planctomycetes and Proteobacteria are abundant in surface ocean metagenomes. Nat Microbiol 3, 804–813, doi:10.1038/s41564-018-0176-9 (2018).

22 Pachiadaki, M. G. et al. Charting the Complexity of the Marine Microbiome through Single-Cell Genomics. Cell 179, 1623–1635 e1611, doi:10.1016/j.cell.2019.11.017 (2019).

23 Paoli, L. et al. Biosynthetic potential of the global ocean microbiome. Nature 607, 111–118, doi:10.1038/s41586-022-04862-3 (2022).

24 Chaffron, S., Rehrauer, H., Pernthaler, J. & von Mering, C. A global network of coexisting microbes from environmental and whole-genome sequence data. Genome Research, doi:10.1101/gr.104521.109 (2010).

25 Machado, D. et al. Polarization of microbial communities between competitive and cooperative metabolism. Nat Ecol Evol 5, 195–203, doi:10.1038/s41559-020-01353-4 (2021).

26 Dal Bello, M., Lee, H., Goyal, A. & Gore, J. Resource-diversity relationships in bacterial communities reflect the network structure of microbial metabolism. Nat Ecol Evol 5, 1424–1434, doi:10.1038/s41559-021-01535-8 (2021).

27 Herold, M. et al. Integration of time-series meta-omics data reveals how microbial ecosystems respond to disturbance. Nat Commun 11, 5281, doi:10.1038/s41467-020-19006-2 (2020).

28 Goyal, A., Wang, T., Dubinkina, V. & Maslov, S. Ecology-guided prediction of cross-feeding interactions in the human gut microbiome. Nat Commun 12, 1335, doi:10.1038/s41467-021-21586-6 (2021).

29 Embree, M., Liu, J. K., Al-Bassam, M. M. & Zengler, K. Networks of energetic and metabolic interactions define dynamics in microbial communities. Proc Natl Acad Sci U S A 112, 15450–15455, doi:10.1073/pnas.1506034112 (2015).

30 Pascual-Garcia, A., Bonhoeffer, S. & Bell, T. Metabolically cohesive microbial consortia and ecosystem functioning. Philos Trans R Soc Lond B Biol Sci 375, 20190245, doi:10.1098/rstb.2019.0245 (2020).

31 Molina, N. & van Nimwegen, E. Scaling laws in functional genome content across prokaryotic clades and lifestyles. Trends Genet 25, 243–247, doi:10.1016/j.tig.2009.04.004 (2009).

32 Mende, D. R. et al. proGenomes: a resource for consistent functional and taxonomic annotations of prokaryotic genomes. Nucleic Acids Res 45, D529–D534, doi:10.1093/nar/gkw989 (2017).

33 Bowers, R. M. et al. Minimum information about a single amplified genome (MISAG) and a metagenome-assembled genome (MIMAG) of bacteria and archaea. Nat Biotechnol 35, 725–731, doi:10.1038/nbt.3893 (2017).

34 Parks, D. H. et al. GTDB: an ongoing census of bacterial and archaeal diversity through a phylogenetically consistent, rank normalized and complete genome-based taxonomy. Nucleic Acids Research 50, D785–D794, doi:10.1093/nar/gkab776 (2021).

35 Sunagawa, S. et al. Ocean plankton. Structure and function of the global ocean microbiome. Science 348, 1261359, doi:10.1126/science.1261359 (2015).

36 van Nimwegen, E. Scaling laws in the functional content of genomes. Trends Genet 19, 479–484, doi:10.1016/S0168-9525(03)00203-8 (2003).

37 Swan, B. K. et al. Prevalent genome streamlining and latitudinal divergence of planktonic bacteria in the surface ocean. Proceedings of the National Academy of Sciences 110, 11463–11468, doi:10.1073/pnas.1304246110 (2013).

38 Romine, M. F., Rodionov, D. A., Maezato, Y., Osterman, A. L. & Nelson, W. C. Underlying mechanisms for syntrophic metabolism of essential enzyme cofactors in microbial communities. ISME J 11, 1434–1446, doi:10.1038/ismej.2017.2 (2017).

39 Zoccarato, L., Sher, D., Miki, T., Segre, D. & Grossart, H. P. A comparative whole-genome approach identifies bacterial traits for marine microbial interactions. Commun Biol 5, 276, doi:10.1038/s42003-022-03184-4 (2022).

40 Sanudo-Wilhelmy, S. A. et al. Multiple B-vitamin depletion in large areas of the coastal ocean. Proc Natl Acad Sci U S A 109, 14041–14045, doi:10.1073/pnas.1208755109 (2012).

41 Salazar, G. et al. Gene Expression Changes and Community Turnover Differentially Shape the Global Ocean Metatranscriptome. Cell 179, 1068–1083 e1021, doi:10.1016/j.cell.2019.10.014 (2019).

42 Krause, E. et al. Small changes in pH have direct effects on marine bacterial community composition: a microcosm approach. PLoS One 7, e47035, doi:10.1371/journal.pone.0047035 (2012).

43 Nelson, K. S., Baltar, F., Lamare, M. D. & Morales, S. E. Ocean acidification affects microbial community and invertebrate settlement on biofilms. Sci Rep 10, 3274, doi:10.1038/s41598-020-60023-4 (2020).

44 Joint, I., Doney, S. C. & Karl, D. M. Will ocean acidification affect marine microbes? ISME J 5, 1–7, doi:10.1038/ismej.2010.79 (2011).

45 Lomas, M. W. et al. Effect of ocean acidification on cyanobacteria in the subtropical North Atlantic. Aquatic Microbial Ecology 66, 211–222 (2012).

46 Browning, T. J. et al. Iron limitation of microbial phosphorus acquisition in the tropical North Atlantic. Nat Commun 8, 15465, doi:10.1038/ncomms15465 (2017).

47 Ustick, L. J. et al. Metagenomic analysis reveals global-scale patterns of ocean nutrient limitation. Science 372, 287–291, doi:10.1126/science.abe6301 (2021).

48 Fuhrman, J. A. et al. Annually reoccurring bacterial communities are predictable from ocean conditions. Proceedings of the National Academy of Sciences of the United States of America 103, 13104–13109, doi:10.1073/pnas.0602399103 (2006).

49 Fuhrman, J. A., Cram, J. A. & Needham, D. M. Marine microbial community dynamics and their ecological interpretation. Nat Rev Microbiol 13, 133–146, doi:10.1038/nrmicro3417 (2015).

50 Tackmann, J., Matias Rodrigues, J. F. & von Mering, C. Rapid Inference of Direct Interactions in Large-Scale Ecological Networks from Heterogeneous Microbial Sequencing Data. Cell Syst 9, 286–296 e288, doi:10.1016/j.cels.2019.08.002 (2019).

51 Giri, S. et al. Metabolic dissimilarity determines the establishment of cross-feeding interactions in bacteria. Curr Biol 31, 5547–5557 e5546, doi:10.1016/j.cub.2021.10.019 (2021).

52 Smillie, C. S. et al. Ecology drives a global network of gene exchange connecting the human microbiome. Nature 480, 241–244, doi:10.1038/nature10571 (2011).

53 Machado, D., Andrejev, S., Tramontano, M. & Patil, K. R. Fast automated reconstruction of genome-scale metabolic models for microbial species and communities. Nucleic Acids Res 46, 7542–7553, doi:10.1093/nar/gky537 (2018).

54 Lieven, C. et al. MEMOTE for standardized genome-scale metabolic model testing. Nat Biotechnol 38, 272–276, doi:10.1038/s41587-020-0446-y (2020).

55 Zelezniak, A. et al. Metabolic dependencies drive species co-occurrence in diverse microbial communities. Proc Natl Acad Sci U S A 112, 6449–6454, doi:10.1073/pnas.1421834112 (2015).

56 D’Souza, G. et al. Ecology and evolution of metabolic cross-feeding interactions in bacteria. Nat Prod Rep 35, 455–488, doi:10.1039/c8np00009c (2018).

57 Hillesland, K. L. & Stahl, D. A. Rapid evolution of stability and productivity at the origin of a microbial mutualism. Proc Natl Acad Sci U S A 107, 2124–2129, doi:10.1073/pnas.0908456107 (2010).

58 Pande, S. et al. Fitness and stability of obligate cross-feeding interactions that emerge upon gene loss in bacteria. The ISME journal, doi:10.1038/ismej.2013.211 (2013).

59 Johnson, W. M. et al. Auxotrophic interactions: a stabilizing attribute of aquatic microbial communities? FEMS Microbiol Ecol 96, doi:10.1093/femsec/fiaa115 (2020).

60 Zhang, H. & Yang, C. Arginine and nitrogen mobilization in cyanobacteria. Mol Microbiol 111, 863–867, doi:10.1111/mmi.14204 (2019).

61 Majumdar, R. et al. Glutamate, Ornithine, Arginine, Proline, and Polyamine Metabolic Interactions: The Pathway Is Regulated at the Post-Transcriptional Level. Front Plant Sci 7, 78, doi:10.3389/fpls.2016.00078 (2016).

62 Grzymski, J. J. & Dussaq, A. M. The significance of nitrogen cost minimization in proteomes of marine microorganisms. ISME J 6, 71–80, doi:10.1038/ismej.2011.72 (2012).

63 Shenhav, L. & Zeevi, D. Resource conservation manifests in the genetic code. Science 370, 683–687, doi:10.1126/science.aaz9642 (2020).

64 Mee, M. T., Collins, J. J., Church, G. M. & Wang, H. H. Syntrophic exchange in synthetic microbial communities. Proceedings of the National Academy of Sciences of the United States of America 111, E2149–2156, doi:10.1073/pnas.1405641111 (2014).

65 Wienhausen, G., Bittner, M. J. & Paerl, R. W. Key Knowledge Gaps to Fill at the Cell-To-Ecosystem Level in Marine B-Vitamin Cycling. Frontiers in Marine Science 9, doi:10.3389/fmars.2022.876726 (2022).

66 Gomez-Consarnau, L. et al. Mosaic patterns of B-vitamin synthesis and utilization in a natural marine microbial community. Environ Microbiol 20, 2809–2823, doi:10.1111/1462-2920.14133 (2018).

67 Paerl, R. W. et al. Prevalent reliance of bacterioplankton on exogenous vitamin B1 and precursor availability. Proc Natl Acad Sci U S A 115, E10447–E10456, doi:10.1073/pnas.1806425115 (2018).

68 Tomas, H., et al. Vitamin interdependencies predicted by metagenomics-informed network analyses validated in microbial community microcosms. bioRxiv, 2023.2001.2027.524772, doi:10.1101/2023.01.27.524772 (2023).

69 Shelton, A. N. et al. Uneven distribution of cobamide biosynthesis and dependence in bacteria predicted by comparative genomics. ISME J 13, 789–804, doi:10.1038/s41396-018-0304-9 (2019).

70 Gruber, K. & Kratky, C. Coenzyme B(12) dependent glutamate mutase. Curr Opin Chem Biol 6, 598–603, doi:10.1016/s1367-5931(02)00368-x (2002).

71 Stewart, K. L., Stewart, A. M. & Bobik, T. A. Prokaryotic Organelles: Bacterial Microcompartments in E. coli and Salmonella. EcoSal Plus 9, doi:10.1128/ecosalplus.ESP-0025-2019 (2020).

72 Morris, J. J., Lenski Richard, E. & Zinser Erik, R. The Black Queen Hypothesis: Evolution of Dependencies through Adaptive Gene Loss. mBio 3, e00036–00012, doi:10.1128/mBio.00036-12 (2012).

73 Mas, A., Jamshidi, S., Lagadeuc, Y., Eveillard, D. & Vandenkoornhuyse, P. Beyond the Black Queen Hypothesis. ISME J 10, 2085–2091, doi:10.1038/ismej.2016.22 (2016).

74 Giovannoni, S. J., Cameron Thrash, J. & Temperton, B. Implications of streamlining theory for microbial ecology. ISME J 8, 1553–1565, doi:10.1038/ismej.2014.60 (2014).

75 Price, M. N. et al. Filling gaps in bacterial amino acid biosynthesis pathways with high-throughput genetics. PLoS Genet 14, e1007147, doi:10.1371/journal.pgen.1007147 (2018).

76 Ferenci, T. Trade-off Mechanisms Shaping the Diversity of Bacteria. Trends Microbiol 24, 209–223, doi:10.1016/j.tim.2015.11.009 (2016).

77 Morris, J. J., Kirkegaard, R., Szul, M. J., Johnson, Z. I. & Zinser, E. R. Facilitation of robust growth of Prochlorococcus colonies and dilute liquid cultures by “helper” heterotrophic bacteria. Applied and environmental microbiology 74, 4530–4534, doi:10.1128/AEM.02479-07 (2008).

78 Jo, C., Bernstein, D. B., Vaisman, N., Frydman, H. M. & Segre, D. Construction and Modeling of a Coculture Microplate for Real-Time Measurement of Microbial Interactions. mSystems, e0001721, doi:10.1128/msystems.00017-21 (2023).

79 Klemetsen, T. et al. The MAR databases: development and implementation of databases specific for marine metagenomics. Nucleic Acids Res 46, D692–D699, doi:10.1093/nar/gkx1036 (2018).

80 Parks, D. H. et al. Recovery of nearly 8,000 metagenome-assembled genomes substantially expands the tree of life. Nat Microbiol 2, 1533–1542, doi:10.1038/s41564-017-0012-7 (2017).

81 Tully, B. J., Sachdeva, R., Graham, E. D. & Heidelberg, J. F. 290 metagenome-assembled genomes from the Mediterranean Sea: a resource for marine microbiology. PeerJ 5, e3558, doi:10.7717/peerj.3558 (2017).

82 Parks, D. H., Imelfort, M., Skennerton, C. T., Hugenholtz, P. & Tyson, G. W. CheckM: assessing the quality of microbial genomes recovered from isolates, single cells, and metagenomes. Genome Res 25, 1043–1055, doi:10.1101/gr.186072.114 (2015).

83 Olm, M. R., Brown, C. T., Brooks, B. & Banfield, J. F. dRep: a tool for fast and accurate genomic comparisons that enables improved genome recovery from metagenomes through de-replication. The ISME Journal 11, 2864–2868, doi:10.1038/ismej.2017.126 (2017).

84 Jain, C., Rodriguez, R. L., Phillippy, A. M., Konstantinidis, K. T. & Aluru, S. High throughput ANI analysis of 90K prokaryotic genomes reveals clear species boundaries. Nat Commun 9, 5114, doi:10.1038/s41467-018-07641-9 (2018).

85 Chaumeil, P. A., Mussig, A. J., Hugenholtz, P. & Parks, D. H. GTDB-Tk: a toolkit to classify genomes with the Genome Taxonomy Database. Bioinformatics 36, 1925–1927, doi:10.1093/bioinformatics/btz848 (2019).

86 Letunic, I. & Bork, P. Interactive Tree Of Life (iTOL) v5: an online tool for phylogenetic tree display and annotation. Nucleic Acids Res 49, W293–W296, doi:10.1093/nar/gkab301 (2021).

87 Zhu, Q. et al. Phylogenomics of 10,575 genomes reveals evolutionary proximity between domains Bacteria and Archaea. Nature Communications 10, 5477, doi:10.1038/s41467-019-13443-4 (2019).

88 Hyatt, D. et al. Prodigal: prokaryotic gene recognition and translation initiation site identification. BMC Bioinformatics 11, 119, doi:10.1186/1471-2105-11-119 (2010).

89 Huerta-Cepas, J. et al. eggNOG 5.0: a hierarchical, functionally and phylogenetically annotated orthology resource based on 5090 organisms and 2502 viruses. Nucleic Acids Res 47, D309–D314, doi:10.1093/nar/gky1085 (2019).

90 Alberti, A. et al. Viral to metazoan marine plankton nucleotide sequences from the Tara Oceans expedition. Sci Data 4, 170093, doi:10.1038/sdata.2017.93 (2017).

91 Langmead, B. & Salzberg, S. L. Fast gapped-read alignment with Bowtie 2. Nature methods 9, 357–359, doi:10.1038/nmeth.1923 (2012).

92 Danecek, P. et al. Twelve years of SAMtools and BCFtools. Gigascience 10, doi:10.1093/gigascience/giab008 (2021).

93 Van Dongen, S. Graph Clustering Via a Discrete Uncoupling Process. SIAM Journal on Matrix Analysis and Applications 30, 121–141, doi:10.1137/040608635 (2008).

